# Improved accuracy of the whole body Center of Mass position through Kalman filtering

**DOI:** 10.1101/2024.07.24.604923

**Authors:** Le Mouel Charlotte

## Abstract

The trajectory of the body center of mass (CoM) is critical for evaluating balance. The position of the CoM can be calculated using either kinematic or kinetic methods. Each of these methods has its limitations, and it is difficult to evaluate their accuracy as there is no ground truth to which the CoM trajectory can be compared. In this paper, we use as ground truth the fact that, during the flight phase of running, the acceleration of the CoM is equal to gravity. We evaluate the accuracy of kinematic models of different complexity and find that the error ranges from 14 % to 36 % of gravity. We propose a novel method for optimally combining kinematic and force plate information. When using this proposed method, the error drops to 1.1 to 3.1 % for all kinematic models. Moreover, the proposed method provides a more reliable evaluation of foot placement control, which is commonly used as an indicator of balance during locomotion.

The code for calculating this optimal combination is available in both Python and Matlab at: https://github.com/charlotte-lemouel/center_of_mass

The documentation is available at: https://center-of-mass.readthedocs.io

## Introduction

To maintain balance during movement, the nervous system must adequately control the forces acting on the body, in particular the weight, acting at the body center of mass (CoM) (Hof et al., 2005; Welch & Ting, 2008; Seethapathi & Srinivasan, 2019). During standing for example, the nervous system contracts lower limb muscles proportionally to the position, velocity and acceleration of the CoM to counteract postural perturbations (Welch & Ting, 2008). During walking, the margin of stability between the position of the extrapolated CoM and the base of support is thought to indicate whether subjects adopt a safe or a risky gait (Hof et al., 2005). During both walking and running on flat terrain, subjects select their next foot placement based on the position and velocity of their CoM (Wang & Srinivasan, 2014; Seethapathi & Srinivasan, 2019). The CoM thus appears to be a critical variable for balance, and its trajectory is often used to evaluate balance and risk of falling.

Unfortunately, the position of the CoM cannot be directly measured. Two types of methods are used to estimate the position of the CoM: kinetic methods which use either the Ground Reaction Force (GRF) or the Center of Pressure (CoP), and kinematic methods which use markers placed on the body. Good quality force-plates have a measurement noise of a few Newton, and can provide a very accurate measure of the CoM acceleration. This acceleration can then be integrated twice to determine position. However, because of the double integration, small offsets in the force measurement result in large drifts in the CoM position. One approach to mitigate this drift is to high-pass filter the force data (Maus et al., 2011). As a result, the kinetic CoM only has high-frequency movements around a constant average position. This approach is therefore only applicable to movements where the person stays in place, such as walking on a treadmill, and cannot capture slow changes of the person’s position within the treadmill, or changes in mean CoM height when transitioning from standing to walking to running (Maus et al., 2011). Another approach to mitigate the drift is to apply the double integration only on short segments such as single steps and re-setting the integration constants at each step (Pavei et al., 2017). This approach is therefore only applicable to short or repetitive movements. Moreover, this method has been shown to be very sensitive to the integration constants used, with a 5 % change in the integration constant of initial velocity resulting in a 10 % change of the antero-posterior excursions of the CoM during a gait cycle in walking (Gutierrez-Farewik et al., 2006). Thus, kinetic methods using the GRF require pre- processing, are very sensitive to the pre-processing used, and cannot capture slow changes in CoM position. An alternative kinetic method is to low-pass filter the CoP (Caron et al., 1997). During quiet standing, this should reflect the low-frequency movements of the CoM. This method is however less accurate than either kinetic methods using the GRF or the kinematic methods described below (Lafond et al., 2004).

An alternative method to determine the CoM position is to use the kinematics. Kinematic methods of varying complexity have been proposed. The simplest use a single marker on the sacrum (Rankin et al., 2014; Seethapathi & Srinivasan, 2019) or markers on the pelvis (Whittle, 1997). In more complex models, the body is subdivided into segments. The CoM of each segment is calculated using marker coordinates and anthropometric tables, and the whole body CoM is determined as the barycentre of the segment CoMs. The most detailed models use a large number of markers to calculate three-dimensional segment coordinate systems, which are then used to determine the position of the segment CoM in the longitudinal, antero- posterior and lateral directions relative to the segment origin (Dumas et al., 2007; Dumas & Wojtusch, 2018). Intermediate models use fewer markers and are only able to locate the segment CoM along the segment’s longitudinal axis (de Leva, 1996; Winter, 2009; Tisserand et al., 2016). Various anthropometric tables and marker sets have been proposed, typically based on photogrammetry of either living subjects (de Leva, 1996; Dumas et al., 2007) or cadavers (Winter, 2009) and in various study populations (Park et al., 1999; Nikolova & Toshev, 2007). Importantly, the choice of kinematic model can have a large influence on the resulting CoM trajectory (Gullstrand et al., 2009; Pavei et al., 2017; Catena et al., 2017). It is however not known which of these models is the most accurate for a given study population. Indeed, since the CoM position cannot be measured directly, no ground truth is available to determine the accuracy of these kinematic models.

Kinetic methods have been proposed as a way of validating the various kinematic models. The most common approach is to compare kinematic and kinetic methods on a single gait cycle (or an average gait cycle). When applied at this short timescale, kinetic methods are assumed to accurately capture the CoM motion. They are therefore used as a reference and the mismatch between kinematic and kinetic methods is thought to reflect mainly the inaccuracy of the kinematic model. When using the pelvis or sacrum as the kinematic estimate, the excursions of the CoM are over-estimated in the vertical and antero-posterior directions relative to the kinetic estimate in able-bodied children and adults (Whittle, 1997; Eames et al., 1999; Gard et al., 2004). The excursions may also be overestimated in the lateral direction and phase-shifted in the antero-posterior direction (Whittle, 1997), although this is not systematically observed (Eames et al., 1999). When using segmental kinematic models, the vertical excursions may also be overestimated (Maus et al., 2011), although this is not systematically found (Eames et al., 1999). During walking in children, the root mean squared (RMS) distance between the kinetic and the segmental kinematic CoM was 15 mm (Gutierrez-Farewik et al., 2006). During walking in adults, the RMS distance ranged from 2.5 mm to 6.7 mm for various segmental models and reached 10.7 mm for the pelvis (Pavei et al., 2017). The RMS was even larger in running, ranging from 4.9 mm to 6.8 mm for segmental models and reaching 14.9 mm for the pelvis (Pavei et al., 2017). There are thus substantial discrepancies between kinematic and kinetic methods during gait, which are largest when using the pelvis or sacrum as the kinematic CoM. It isn’t clear whether these discrepancies entirely reflect the inaccuracy of the kinematic CoM, or whether they also depend on the pre-processing used in the kinetic method.

An alternative way to use kinetics to validate kinematic models is to compare the GRF to the kinematic acceleration obtained by double differentiation of the kinematic CoM. Whereas comparing CoM positions obtained from kinematic and kinetic methods requires pre- processing the GRF, comparing the accelerations does not. Instead, it requires additional pre- processing of the kinematics. Indeed, the double differentiation exacerbates measurement noise, therefore kinematic CoM trajectories are first low-pass filtered (Maus et al., 2011; Simonetti et al., 2021). Although this approach has not been systematically used to compare kinematic models, it has shown discrepancies between kinetic and kinematic accelerations on the order of 10 % of the subject’s weight (Simonetti et al., 2021). Similar discrepancies of at least 10 % of the subject’s weight have been reported when applying inverse kinematics, a method which also relies on anthropometric tables (Hamner et al., 2010; Futamure et al., 2017; Sturdy et al., 2022).

Finally, kinetics can be used to obtain individualised body segment inertial parameters for a personalised kinematic model (Cotton et al., 2008; Jovic et al., 2016; Bonnet et al., 2016; Chebel & Tunc, 2023). This requires an identification phase, during which the subject performs a series of poses. The inertial parameters providing the best fit between the kinematic CoM and the kinetics are determined. This can reduce the mismatch between kinematics and kinetics by 10 to 45 %, but at the cost of the subject performing an elaborate seven minute “dance” during the identification phase (Bonnet et al., 2016). In robotics, the statically equivalent serial chain technique is used to obtain personalised kinematic models (Cotton et al., 2008). This allows the CoM to be obtained from a more limited set of parameters than the whole set of body inertial parameters. The identification these parameters nevertheless requires the subject to perform 75 static poses, and the mismatch between the horizontal CoM and CoP locations during these static poses is still 25.5 mm with the optimal parameters (Chebel & Tunc, 2023). Identification techniques therefore require the subject to perform a long series of poses (which may not be feasible for pathological populations) for only a moderate improvement in accuracy.

To mitigate discrepancies between kinematics and kinetics, a heuristic method combining low-frequency kinematic and high-frequency kinetic information was previously proposed (Maus et al., 2011). This relied on a sigmoid weighting of the Fourier coefficients of the CoM velocities obtained from the kinematics and from the kinetics. The cut-off frequency was determined heuristically based on simulations of human walking with wobbling masses. Unfortunately, the accuracy of this method was not evaluated on real human data.

In this paper, we first propose a method to validate kinematic models that requires neither GRF pre-processing nor low-pass filtering of the kinematics. For this, we use the fact that, during the flight phase of running, when the body is no longer in contact with the ground, the acceleration of the CoM is equal to gravity (i.e. downwards, of amplitude 9.81 m/s^2^). We calculate the kinematic CoM acceleration without applying low-pass filtering. For this, we use data with a sufficient number of flight phases to obtain accurate estimates of the median kinematic CoM acceleration during flight. We compare this kinematic acceleration to the ground truth acceleration to determine the accuracy of the kinematic models.

Second, we propose a novel method which uses a Kalman filter to optimally combine kinematic and kinetic measurements. This method does not require pre-processing of either the GRF or the kinematics. We show that this method substantially improves the accuracy of the reconstructed CoM position. The code for implementing this method is publicly available both in Python and Matlab.

Finally, we determine the influence of CoM measurement accuracy on a commonly used indicator of balance: the control of foot placement according to the CoM state (Seethapathi & Srinivasan, 2019).

## Methods

### Measurement protocol

The data was obtained from two publicly available databases.

**Dataset 1** (Wojtusch & von Stryk, 2015). Two subjects (one male aged 32 years old, one female aged 27 years old) performed various tasks on an instrumented treadmill. We analysed two tasks: running for one minute at 3 m/s and at 4 m/s. For each trial, the recording starts with the treadmill unloaded. After ten seconds, the subject steps onto the treadmill and stands still for ten seconds. The subject then performs the task, then stands still for another ten seconds, steps off the treadmill, and the recording stops ten seconds after the subject has stepped off. Ground reaction forces on the treadmill were recorded at 1000 Hz. The positions of 35 markers placed on anatomical landmarks were recorded at 500 Hz. Further information on the measurement protocol can be found in the database’s documentation available at: https://web.sim.informatik.tu-darmstadt.de/download/ds/HuMoD/Documentation/Documentation.pdf

**Dataset 2** (Srinivasan & Seethapathi, 2019). Eight subjects (three female and five male, age 25 ± 5 years) ran on a treadmill at 2.5, 2.7 and 2.9 m/s for around 3.5 minutes at each speed. For each trial, the recording starts and ends with the person running at steady-state speed. Ground reaction forces on the treadmill were recorded at 1000 Hz. The positions of markers on the lower limbs, including 4 markers on the hip (“roughly at the sacral level”) and a marker on the heel, were recorded at 100 Hz. Further information on the measurement protocol can be found in the corresponding paper (Seethapathi & Srinivasan, 2019).

### Data pre-processing

**Dataset 1**. Over all subjects and trials, 8 markers were invisible for a duration of up to 3 seconds. These gaps were filled through linear interpolation. The raw force data was synchronised to the kinematic data. The drift in the force measurement was estimated for each trial by fitting an affine function to the force recordings during the initial and final ten seconds during which the treadmill was not loaded. This drift was then subtracted to the measured force.

**Dataset 2.** The drift in the force measurement was estimated for each trial by fitting an affine function to the force recordings when the vertical force was below 30 Newton. This drift was then subtracted to the measured force.

### Kinematic Center of Mass

**Dataset 1.** The CoM position was calculated using three kinematic models of decreasing complexity.

The “3D-segments” model is based on the state-of-the-art model proposed by Dumas and colleagues (Dumas et al., 2007; Dumas & Wojtusch, 2018) and presented in detail in Appendix A.

The body is divided into sixteen segments (Appendix A Figure 1). The coordinates of 37 markers are used to calculate three-dimensional segment coordinate systems. An anthropometric table (Dumas et al., 2007; Dumas & Wojtusch, 2018) is then used to place the segment CoM in the longitudinal, antero-posterior and lateral directions relative to the segment origin, and to determine the fraction of the segment’s weight relative to the person’s weight. The whole body CoM is then obtained as a weighted sum of the segment CoMs. The method for calculating segment coordinate systems was modified to accommodate for the specific markers present in the database. Specifically, to calculate the 3D orientation of the forearm and hand segment coordinate systems, the state-of-the-art method uses two markers on each elbow, wrist and hand. However, in the database, only a single marker on each elbow and wrist and no markers on the hands were present. We could therefore only calculate the longitudinal axis of the forearm and hand coordinate systems (Tisserand et al., 2016). We expect the resulting “3D- segments” CoM to be very similar when using the full marker set, as each hand only accounts for 0.5 % of the body weight, and the displacement of the forearm CoM in the antero-posterior and lateral directions is less than 3 % of the forearm’s length. Additionally, to calculate the head coordinate system, the original method uses a marker on the vertex and one on the sellion. Instead, the head coordinate system was calculated from markers on the left and right tragion (see Appendix A II.3.b.).

**Figure 1.**
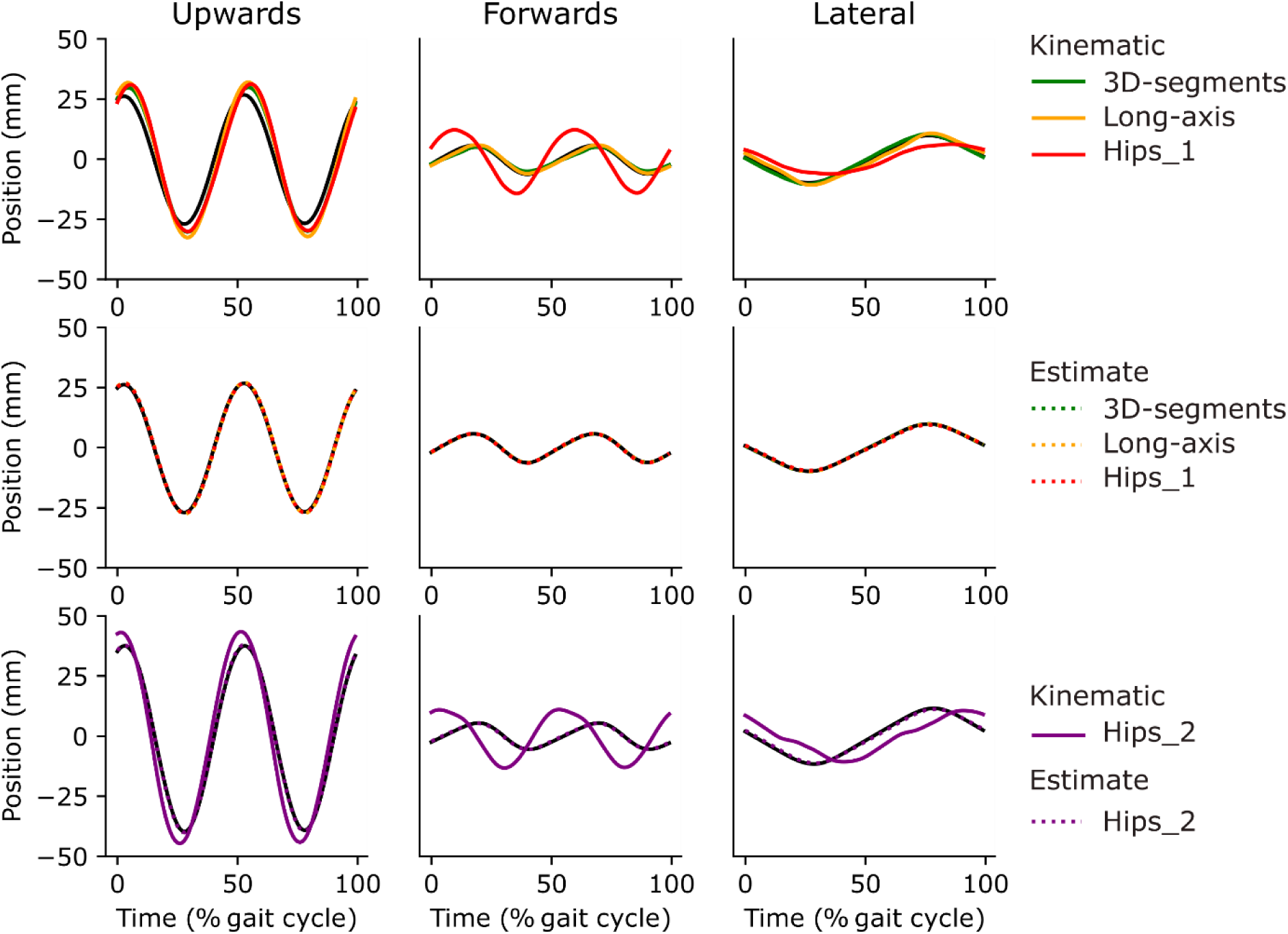
Periodic positions. Kinetic periodic positions (black), kinematic periodic positions (full lines) for the 3D-segments (green), Long-axis (yellow) Hips_1 (red) and Hips_2 (purple) models, and estimate periodic positions (dashed lines) obtained when applying the Kalman filter to each kinematic model with the optimal gains, in the upwards (left column), forwards (middle column) and lateral (right column) directions, averaged over subjects and trials as a % percent of the gait cycle (with flight onset occurring at 50 % of the gait cycle). The first two rows correspond to dataset 1 and the third row to dataset 2. The periodic positions for each subject and trial of dataset 1 are shown in Supplementary Figure 1.

The “Long-axis” model corresponds to the method proposed by Tisserand and colleagues (Tisserand et al., 2016) which is a simplified version of the state-of-the-art method, requiring only 13 markers. The body is divided into nine segments (each foot is fused with the shank, each hand is fused with the forearm, and the pelvis, abdomen, thorax and head are fused into a single trunk segment). This only allows for the calculation of the longitudinal axis for each segment coordinate system. The segment CoMs and weights are then obtained in the same way as for the “3D-segments” model. The full description of the 3D-segments and Long-axis models is provided in Appendix A.

The “Hips_1” model is the average position of four markers placed on the left and right anterior and posterior illiac spines.

**Dataset 2**. The “Hips_2” model is the average position of four markers placed “roughly at the sacral level” (Seethapathi & Srinivasan, 2019).

### Acceleration during the flight phase of running

The onset of the flight (respectively stance) phases was determined as the moment when the vertical ground reaction force dropped below (respectively rose above) 1 % of the subject’s weight. For dataset 1, each subject’s weight was calculated as the median vertical ground reaction force during the initial and final quiet standing periods of the subject’s two trials. The first ten and last five steps were removed from the analysis to ensure the subject was in steady-state running. For dataset 2, each subject’s weight was calculated as the mean across the subject’s three trials of the vertical force from the first stance onset to the last stance onset of each trial.

The kinematic CoM accelerations were obtained by applying a Savitsky-Golay filter to the kinematic CoM positions. The filter was a second order polynomial fit on a sliding window of 3 samples. The lateral acceleration was flipped for all flight phases preceding a right stance phase. The median CoM acceleration during flight was calculated, and the ground truth acceleration (downwards of amplitude 9.81 m/s^2^) was subtracted to the median acceleration to obtain the Acceleration Bias in each direction. The one sample sign test (which is a non-parametric test) was used to determine whether this Acceleration Bias was significantly different from zero in each direction for each kinematic model.

We obtained a 95 % confidence interval on the Acceleration Bias using the bootstrapping method described by Ott and Longnecker (Ott & Longnecker, 2016) in section 5.9. Specifically, let *N* be the number of samples. Each sample has a 50% chance of being either above or below the median. We sort the samples according to their value. Let the *n*^th^ sample correspond to the lower bound of the 95 % confidence interval. This means that only (*n-1*) of the *N* samples are below the median. We use the percent point function of the binomial distribution with parameters *N* and 0.5 to determine *n* and thus the lower bound. We use the same approach to determine the upper bound of the 95 % confidence interval.

Finally, the Acceleration Bias Amplitude was calculated as the root of the sum of the squares of the biases in the upwards, forwards and lateral directions.

### Discrepancy between kinematic and kinetic positions over a gait cycle

In addition to comparing the kinematic acceleration to the ground truth during flight, we also performed a more classical analysis, comparing the kinematic and kinetic CoM positions over a gait cycle. As mentioned in the introduction, calculating the CoM position from double integration of the force is highly sensitive to the integration constants used. To mitigate this problem, the CoM trajectories were calculated over an average gait cycle rather than a single gait cycle, and the integration constants were chosen so as to impose periodicity of the average trajectory. First, kinematic data was upsampled to the kinematic frequency (1000 Hz) through linear interpolation. Then, for each subject and running speed, we averaged the kinematic CoM positions and kinetic CoM acceleration over all gait cycles. We flipped the position and acceleration in the lateral direction for all gait cycles starting with a left stance. We then averaged together left and right gait cycles. We then subtracted an affine function of time from the kinematic position to obtain a periodic signal. For the kinetics, the mean was subtracted from the kinetic acceleration, and this zero-mean acceleration was integrated to obtain a periodic velocity signal. The mean was then subtracted from this velocity, and this zero-mean velocity was integrated to obtain a periodic kinetic position signal. Finally, the mean position was subtracted for both the kinematic and the kinetic periodic positions.

For each dimension, we calculated the squared difference between the kinematic and kinetic periodic positions, averaged this over the duration of the gait cycle, over subjects and over trials, and reported the root of this mean as the Position Error. This Position Error corresponds to the standard deviation of the difference between the kinematics and kinetics. We therefore calculated a 95 % confidence interval on the Position Error using the method described by Ott and Longnecker (Ott and Longnecker, 2016) in section 7.2. Let N be the number of samples and *χ*^2^_0.025_ and *χ*^2^_0.975_ be the percent point functions of the chi-square distribution with N-1 degrees of freedom evaluated at 2.5 % and 97.5%. Then the lower and upper bounds are equal to the Position Error multiplied respectively by 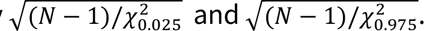.

Additionally, we calculated the (3-dimensional) distance between the kinematic and kinetic periodic positions. The Mann-Whitney U test (which is a non-parametric test) was used to determine whether this distance was significantly lower for the 3D-segments compared to the Long-axis and Hips_1 kinematic models, and for the Long-axis compared to the Hips_1 models. Finally, the square of this distance was averaged aver duration, subjects and trials, and the root of this mean is reported as the Position Error Amplitude. For comparison, for each dataset, the square of the amplitude of the kinetic periodic positions was averaged aver duration, subjects and trials, and the root of this mean is reported as the Kinetic Amplitude.

### Optimal combination of force and kinematic measurements

To improve the accuracy of the CoM measurement, we used a Kalman filter to optimally combine kinematic and kinetic information (with kinematic data upsampled to the kinetic frequency through linear interpolation), presented in detail in Appendix B.

In this framework, we model the inaccuracy in the kinematic CoM position and in the kinetic CoM acceleration as additive white Gaussian noise with standard deviation *p*_*std*_ (in meters) for position and *a*_*std*_ (in m/s^2^) for acceleration. The filter’s steady-state error, gains and transfer function (derived in Appendix B) depend only on the noise ratio *r* of position noise to acceleration noise, normalized by the sampling frequency F (in Hertz):

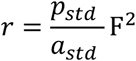

The acceleration noise *a*_*std*_ was obtained by dividing the force noise *f*_*std*_ by the subject’s mass. The force noise *f*_*std*_ in each direction was taken as the standard deviation of the force measurement in dataset 1 when the force plates were empty (after pre-processing the force signal to remove drift, as described above). The position noise *p*_*std*_ cannot be measured directly. We varied *p*_*std*_ from 0.2 mm to 20 mm. For each value of *p*_*std*_, the optimal gains were derived and the CoM estimate was obtained by applying the Kalman filter with these gains. The Acceleration Bias in each direction during flight, confidence interval of the Acceleration Bias and Total Acceleration Bias were calculated for the estimate in the same way as for the kinematic CoM, described above. The estimate periodic position, Position Error in each direction, confidence interval of the Position Error and the Position Error Amplitude were also calculated for the estimate in the same way as for the kinematic CoM.

For each value of *p*_*std*_, the error in the combined CoM due to kinetic noise was estimated by applying the corresponding Kalman filter to the force recordings in dataset 1 during the initial and final ten seconds of each trial during which the treadmill was not loaded. The Kinetic Error Amplitude was taken as the RMS amplitude of the resulting position. A 95 % confidence interval on the Kinetic Error in each direction was obtained by multiplying the Kinetic Error by 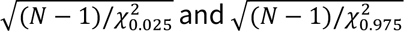 for the lower and upper bounds.

The total estimation error is due to both the kinematic and the kinetic noise. Considering that these two sources of noise are independent, the variance of the estimator error is assumed to be the sum of the variances due to kinematic and kinetic noise. The total estimation error was thus taken as the root of the sum of the squared Kinetic Error Amplitude and the squared Position Error Amplitude. The optimal *p*^∗^ minimizing this total error was determined. The Mann-Whitney U test was used to determine whether the Position Error Amplitude was significantly reduced when applying the Kalman filter with this optimal *p*^∗^ .

### Lateral foot placement predictor

During running, the lateral foot placement at the onset of stance can be predicted from the state of the CoM at the preceding flight apex: the foot is thus placed more laterally when the lateral CoM velocity during flight is larger (Seethapathi & Srinivasan, 2019). Following the approach in Seethapathi & Srinivasan (2019), the lateral foot placement was taken as the lateral distance between the heel marker and the CoM at stance onset. The CoM apex was taken as the time during the flight phase when the vertical position of the CoM was highest. The CoM state was composed of the CoM lateral velocity, forwards velocity and height. A linear regression between the lateral foot placement and the CoM state at the preceding apex was performed for each subject, running speed, leg and CoM measurement (i.e. for the estimate and kinematic CoM, and for each of the kinematic models separately). For each CoM measurement, the median r-square of the regression was reported. The 95% confidence interval on the median r-square was obtained using the same method as for the Acceleration Bias. The Mann-Whitney U test was used to determine wether the r-square was significantly increased when applying the Kalman filter with the optimal *p*^∗^ .

## Results

### (In-)Accuracy of the kinematic models

All four kinematic models showed a significant Acceleration Bias during the flight phase of running in the upwards and forwards directions, and the Hips_1 and Hips_2 models additionally had a significant bias in the lateral direction (p < 0.001 one sample sign test, Table 1). The downwards acceleration was larger than gravity, indicating that all four kinematic models exaggerate the vertical motion of the CoM. The amplitude of the Acceleration Bias Amplitude was 1.3 m/s^2^ for the 3D-segments model, 2.3 m/s^2^ for the Long-axis model, 3.6 m/s^2^ for the Hips_1 and 2.6 m/s^2^ for the Hips_2. The state-of-the-art kinematic model thus has a 14 % error in acceleration during the flight phase of running, and this error increases to 37 % for the most reduced model.

**Table 1.** Acceleration Bias. Acceleration Bias in each direction and Acceleration Bias Amplitude, in m/s^2^, for the four kinematic models and the estimates obtained when applying the Kalman filter to each kinematic model with the optimal gains. The 95 % confidence interval is indicated for the Acceleration Bias, and the Acceleration Bias Amplitude is additionally reported as a % of gravity.

|  |  |  | Upwards | Forwards | Lateral | Amplitude |
| --- | --- | --- | --- | --- | --- | --- |
| **Kinematic** | Dataset 1 | 3D-segments | -1.17<br>(-1.28 to -1.04) | 0.66<br>(0.49 to 0.81) | -0.05<br>(-0.18 to 0.08) | 1.34<br>(13.7 %) |
|  |  | Long-axis | -2.20<br>(-2.31 to -2.10) | 0.61<br>(0.46 to 0.76) | -0.12<br>(-0.25 to 0.01) | 2.29<br>(23.3 %) |
|  |  | Hips_1 | -2.32<br>(-2.47 to -2.18) | -2.65<br>(-2.80 to -2.50) | -0.50<br>(-0.62 to -0.38) | 3.56<br>(36.3 %) |
|  | Dataset 2 | Hips_2 | -1.46<br>(-1.47 to -1.44) | -2.15<br>(-2.18 to -2.13) | -0.25<br>(-0.26 to -0.24) | 2.61<br>(26.6 %) |
| **Estimate** | Dataset 1 | 3D-segments | -0.20<br>(-0.20 to -0.20) | -0.12<br>(-0.12 to -0.11) | 0.00<br>(-0.00 to 0.00) | 0.23<br>(2.3 %) |
|  |  | Long-axis | -0.25<br>(-0.25 to -0.25) | -0.12<br>(-0.12 to -0.12) | -0.03<br>(-0.03 to -0.03) | 0.28<br>(2.8 %) |
|  |  | Hips_1 | -0.19<br>(-0.19 to -0.18) | -0.24<br>(-0.24 to -0.23) | -0.04<br>(-0.04 to -0.04) | 0.30<br>(3.1 %) |
|  | Dataset 2 | Hips_2 | -0.06<br>(-0.06 to -0.06) | -0.07<br>(-0.07 to -0.07) | -0.04<br>(-0.04 to -0.04) | 0.10<br>(1.1 %) |

The error in the kinematic CoM position cannot be measured directly. Instead, we estimated the error over a gait cycle by comparing the kinematic periodic position (Figure 1.A, green for 3D-segments, yellow for Long-axis and red for Hips_1; and Figure 1.C purple for Hips_2) to the reference kinetic periodic position (Figure 1, black). All four kinematic models exaggerated the vertical oscillations of the CoM (Figure 1.A, C left). Moreover, the hips movement in the horizontal plane does not reflect the movement of the kinetic CoM, with exaggerated antero- posterior motion (Figure 1.A, C middle, red and purple compared to black) and phase-shifted lateral motion (Figure 1.A, C right, red and purple compared to black). The Position Error for all models is indicated in Table 2. The Position Error Amplitude was significantly larger for the Hips_1 (12.0 mm) followed by the Long-axis model (6.9 mm) then the 3D-segments model (5.9 mm) (p < 0.001 Mann-Whitney U test). The Position Error Amplitude was 15.4 mm for the Hips_2. The average amplitude of the kinetic CoM motion (Kinetic Amplitude) was 21.4 mm for dataset 1 and 31.4 mm for dataset 2. The state-of-the-art kinematic model thus has a 27 % error in position during running, and this error increases to 56 % for the most reduced model.

**Table 2.** Position Error. Position Error in each direction and Position Error Amplitude (in mm) for the four kinematic models and the estimates obtained when applying the Kalman filter to each kinematic model with the optimal gains. The 95 % confidence interval is indicated for Position Error, and the Position Error Amplitude is additionally reported as a % of the Kinetic Amplitude (21.4 mm for dataset 1, 31.4 mm for dataset 2).

|  |  |  | Upwards | Forwards | Lateral | Amplitude |
| --- | --- | --- | --- | --- | --- | --- |
| **Kinematic** | Dataset 1 | 3D-segments | 5.55<br>(5.40 to 5.71) | 1.12<br>(1.09 to 1.15) | 0.99<br>(0.96 to 1.01) | 5.75<br>(26.8 %) |
|  |  | Long-axis | 6.49<br>(6.31 to 6.67) | 0.87<br>(0.85 to 0.90) | 1.75<br>(1.70 to 1.80) | 6.78<br>(31.6 %) |
|  |  | Hips_1 | 7.43<br>(7.22 to 7.63) | 7.95<br>(7.73 to 8.17) | 4.95<br>(4.82 to 5.09) | 11.95<br>(55.7 %) |
|  | Dataset 2 | Hips_2 | 8.26<br>(8.18 to 8.35) | 10.07<br>(9.97 to 10.18) | 8.27<br>(8.19 to 8.36) | 15.43<br>(49.1 %) |
| **Estimate** | Dataset 1 | 3D-segments | 0.55<br>(0.54 to 0.57) | 0.04<br>(0.03 to 0.04) | 0.10<br>(0.10 to 0.11) | 0.56<br>(2.6 %) |
|  |  | Long-axis | 0.60<br>(0.58 to 0.61) | 0.03<br>(0.03 to 0.03) | 0.17<br>(0.16 to 0.17) | 0.62<br>(2.9 %) |
|  |  | Hips_1 | 0.52<br>(0.50 to 0.53) | 0.22<br>(0.22 to 0.23) | 0.40<br>(0.39 to 0.41) | 0.69<br>(3.2 %) |
|  | Dataset 2 | Hips_2 | 0.62<br>(0.61 to 0.63) | 0.26<br>(0.25 to 0.26) | 0.67<br>(0.66 to 0.67) | 0.95<br>(3.0 %) |

### CoM estimator combining kinematic and kinetic information

To improve the accuracy of the CoM measurement, we used a Kalman filter to optimally combine kinematic and kinetic information. We found that this optimal integration depends only on the noise ratio *r* of position noise *p*_*std*_ to acceleration noise *a*_*std*_, normalized by the sampling frequency F (in Hertz):

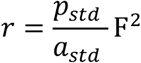

The amplitude of the transfer function for position and acceleration measurements is shown in Figure 2 for different values of this ratio. When the ratio is small (purple curves in Figure 2), the optimal integration relies mainly on position information. This is coherent with the fact that a small ratio indicates that position measurements are far more accurate than acceleration measurements (*p*_*std*_F^2^ ≪ *a*_*std*_). When the ratio is large (yellow curves in Figure 2), the optimal integration relies mainly on acceleration information, except at low frequencies where it relies on position information. Indeed, in this case, despite acceleration measurements being far more accurate than position measurements (*p*_*std*_F^2^ ≫ *a*_*std*_), the slow fluctuations in CoM position cannot be accurately determined from double integration of the acceleration. The ratio determines the cutoff frequency for position and acceleration.

**Figure 2.**
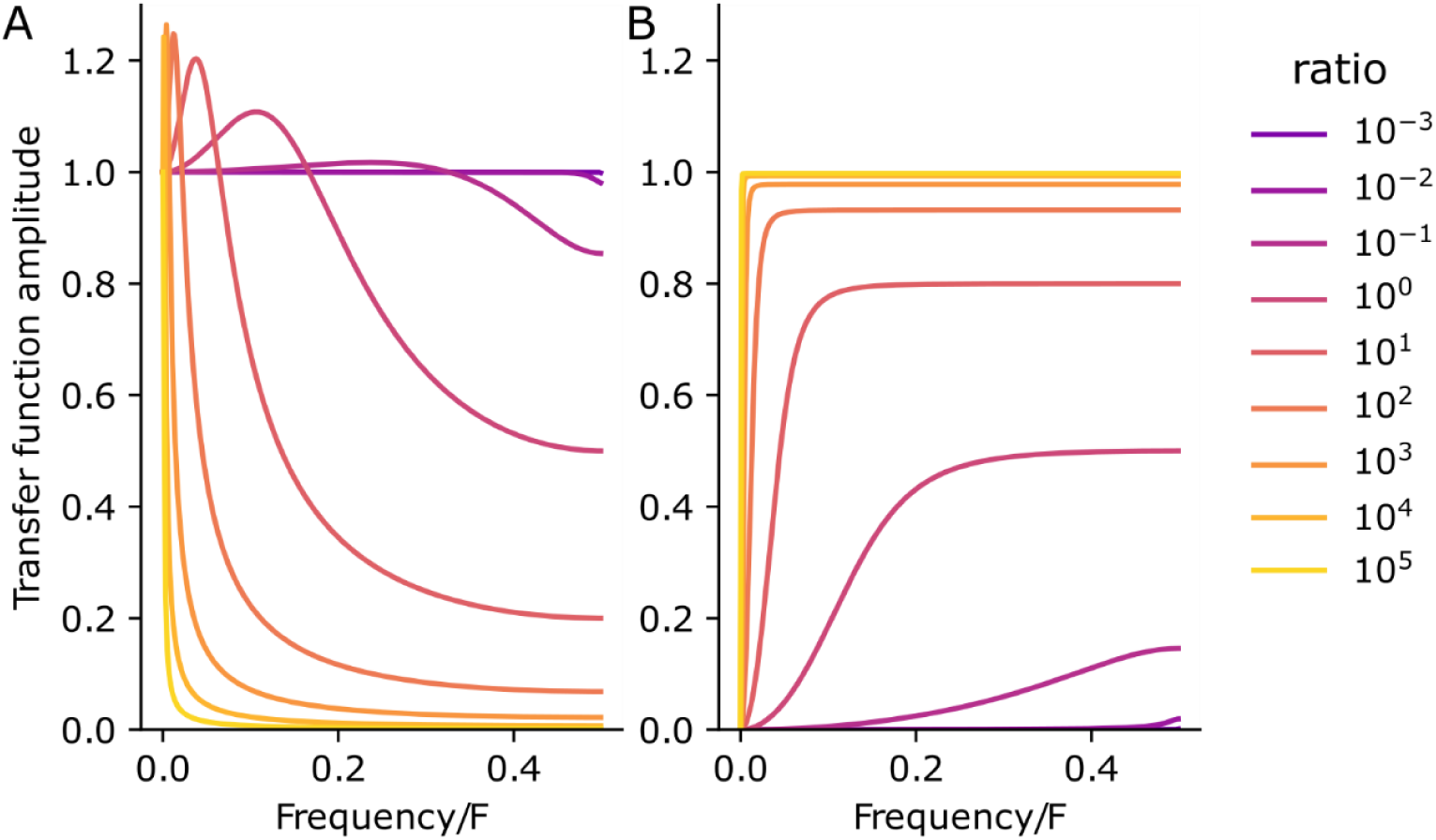
Transfer function. Amplitude of the transfer function from the kinematic to the estimate position (A), and from the double integral of acceleration to the estimate position (B), as a function of the frequency divided by the sampling frequency *F*, for different values of the noise ratio *r*.

### Improved accuracy of the estimate

The position of the CoM was estimated by combining each of the kinematic models with kinetic information using the proposed Kalman filter. Since the parameter *p*_*std*_ could not be determined a priori, the estimation was performed for *p*_*std*_ ranging from 0.2 to 20 mm. With increasing *p*_*std*_, there was a dramatic reduction in both the Acceleration Bias (Figure 3.A) and the Position Error (Figure 3.B) in all three directions for all four models. There was also a gradual increase in the Kinetic Error with *p*_*std*_ (Figure 3.B in black). Note that this Kinetic Error does not seem to increase with time, and is much smaller than the drift in position when the CoM is obtained by double integration of the kinetic signal (Supplementary Figure 2). Adjusting the parameter *p*_*std*_ thus determines a trade-off between the amount of kinematic and kinetic noise in the estimation. For the 3D-segments model, the total estimation error was minimal for *p*_*std*_ = 1.8 *mm* with a Kinetic Error Amplitude of 0.65 mm. For the Long-axis model, it was minimal for *p*_*std*_ = 2.0 *mm* with a Kinetic Error Amplitude of 0.71 mm. For the Hips_1 model, it was minimal for *p*_*std*_ = 2.5 *mm* with a Kinetic Error Amplitude of 0.84 mm. For the Hips_2 model, it was minimal for *p*_*std*_ = 3.9 *mm* with a Kinetic Error Amplitude of 1.15 mm. Combining kinematic and kinetic information thus led to a drastic reduction in kinematic error, with only a small addition of kinetic noise (< 1.2 mm for all models).

**Figure 3.**
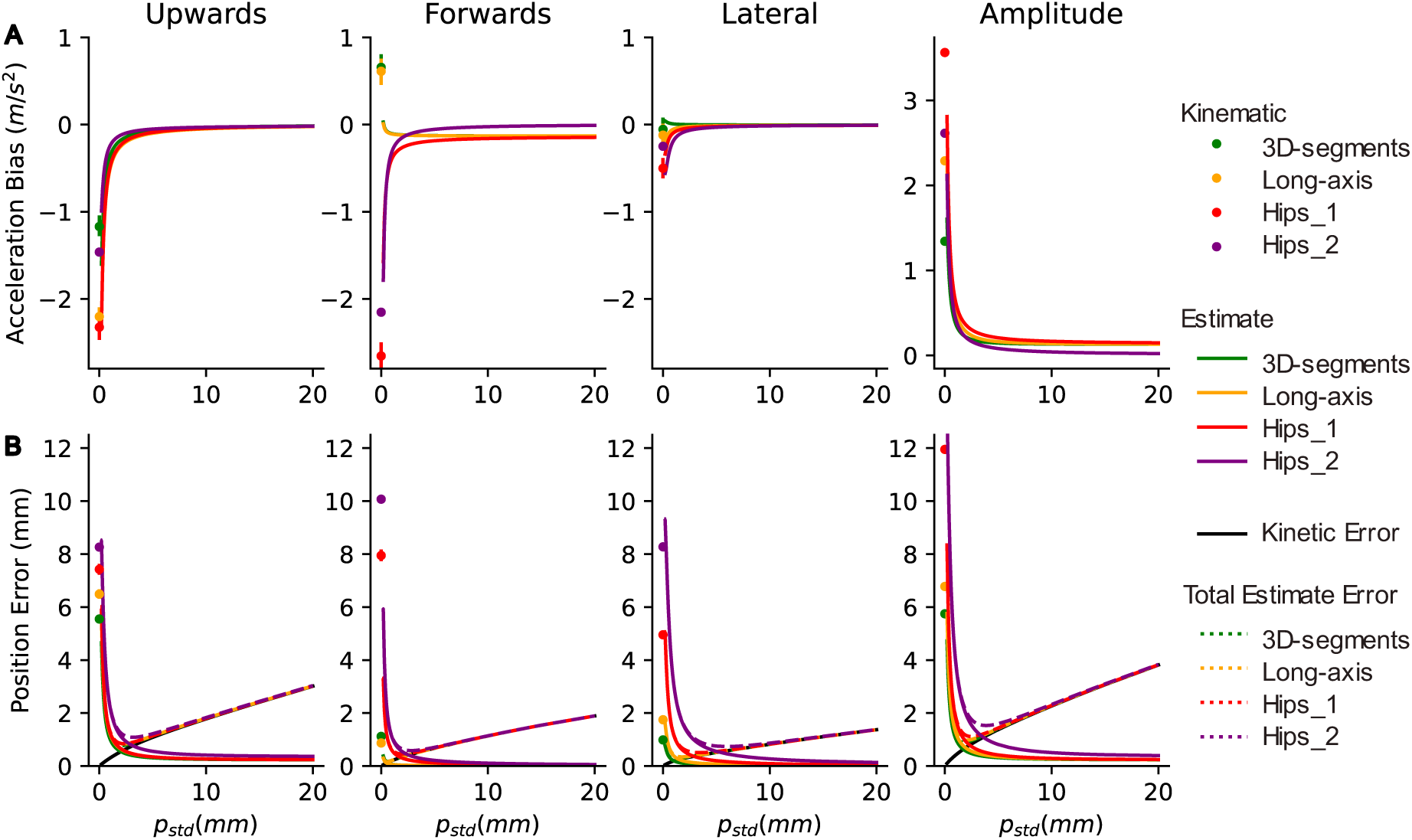
Improved accuracy of the estimate. (A) Acceleration Bias in each direction and Acceleration Bias Amplitude for the 3D-segments (green), Long-axis (yellow), Hips_1 (red) and Hips_2 (purple) models. The values for the kinematic CoM are indicated as a dot (at *p*_*std*_ = 0) with a bar indicating the 95 % confidence interval. The values for the estimate CoM are indicated as a full line over the range of *p*_*std*_, whose thickness corresponds to the 95 % confidence interval (note that this confidence interval is very narrow) . (B) Position Error for the kinematic CoM (dot at *p*_*std*_ = 0 with a bar indicating the 95 % confidence interval) and the estimate CoM (full colored lines, whose thickness corresponds to the 95 % confidence interval), Kinetic Error (black) and total estimation error (dashed colored lines).

When using the optimal *p*_*std*_, the estimate was far more accurate than the kinematic CoM. The Acceleration Bias Amplitude decreased from 13.7 % to 2.3 % of gravity for the 3D-segments model, from 23.3 % to 2.8 % for the Long-axis model; from 36.3 % to 3.1 % for the Hips_1 and from 26.6 % to 1.1 % for the Hips_2 (Table 1). The estimate periodic positions (Figure 1.B, C dashed lines) match the kinetic periodic positions (Figure 1, black lines) much more closely than the kinematic periodic positions for all four kinematic models (Figure 1.A, C). For the 3D- segments model, the Position Error Amplitude significantly decreased (p < 0.001, Mann-Whitney U test, Table 2) from 5.8 mm to 0.6 mm, with a total estimation error of 0.9 mm (4.2 % of the Kinetic Amplitude). For the Long-axis model, it significantly decreased from 6.8 mm to 0.6, with a total estimation error of 1.0 mm (4.6 % of the Kinetic Amplitude). For the Hips_1, it significantly decreased from 12.0 mm to 0.7 mm, with a total estimation error of 1.1 mm (5.2 % of the Kinetic Amplitude). For the Hips_2, it significantly decreased from 15.4 mm to 1.0 mm, with a total estimation error of 1.5 mm (4.9 % of the Kinetic Amplitude). The optimal estimation error was thus 6 to 10 times smaller than the kinematic error for position and 5 to 10 times smaller for acceleration.

Large improvements in accuracy can be obtained for a wide range of the parameter *p*_*std*_ (Figure 4). An acceleration error of less than 0.6 m/s^2^ can be obtained for all models when the parameter is at least half its optimal value (Figure 4.A). A position error of less than 3.4 mm can be obtained when the parameter is between a third and three times its optimal value (Figure 4.B). Drastic decreases in the error can thus be obtained without requiring the optimal *p*_*std*_ to be accurately determined.

**Figure 4.**
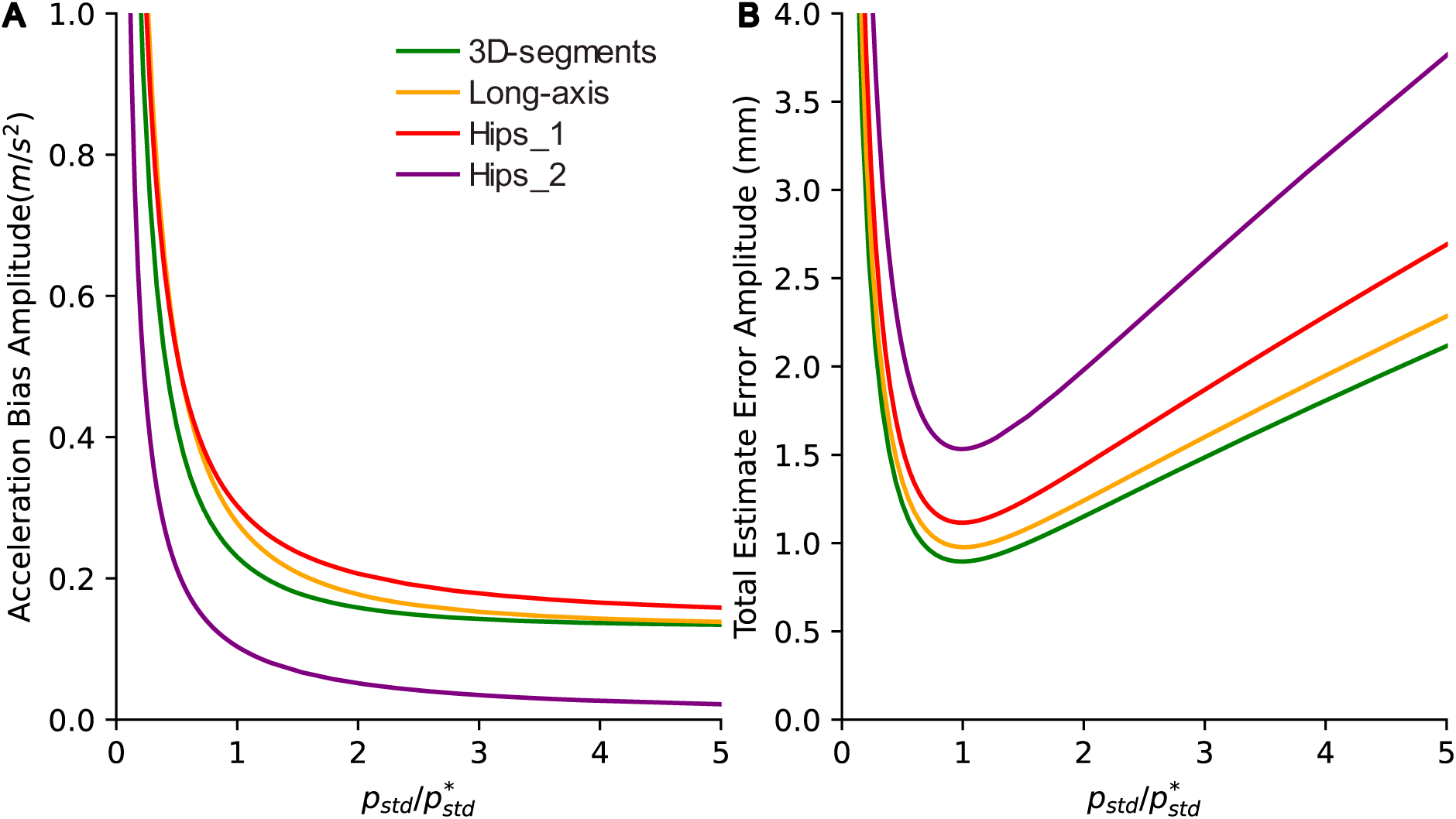
Sensitivity to *p*_*std*_. Amplitude of the estimate Acceleration Bias (A) and total estimate error (B) for the 3D-segments (green), Long-axis (yellow), Hips_1 (red) and Hips_2 (purple) models, as a function of the ratio of *p*_*std*_ to the optimal *p*^∗^_*std*_.

### Improved accuracy of the lateral foot placement prediction

The lateral position of the foot relative to the CoM at the stance onset was predicted from the CoM state at the preceding flight apex. For all four kinematic models, the prediction was much more accurate when it was based on the estimate rather than the kinematic CoM state (Table 3, Figure 5). When using the optimal *p*_*std*_, the median r-square of the linear regression was significantly increased (p < 0.001, Mann-Whitney U test) from 0.38 for the kinematics to 0.58 for the estimate for the 3D-segments, from 0.33 to 0.58 for the Long-axis, from 0.13 to 0.58 for the Hips_1 and from 0.23 to 0.52 for the Hips_2 models.

**Figure 5.**
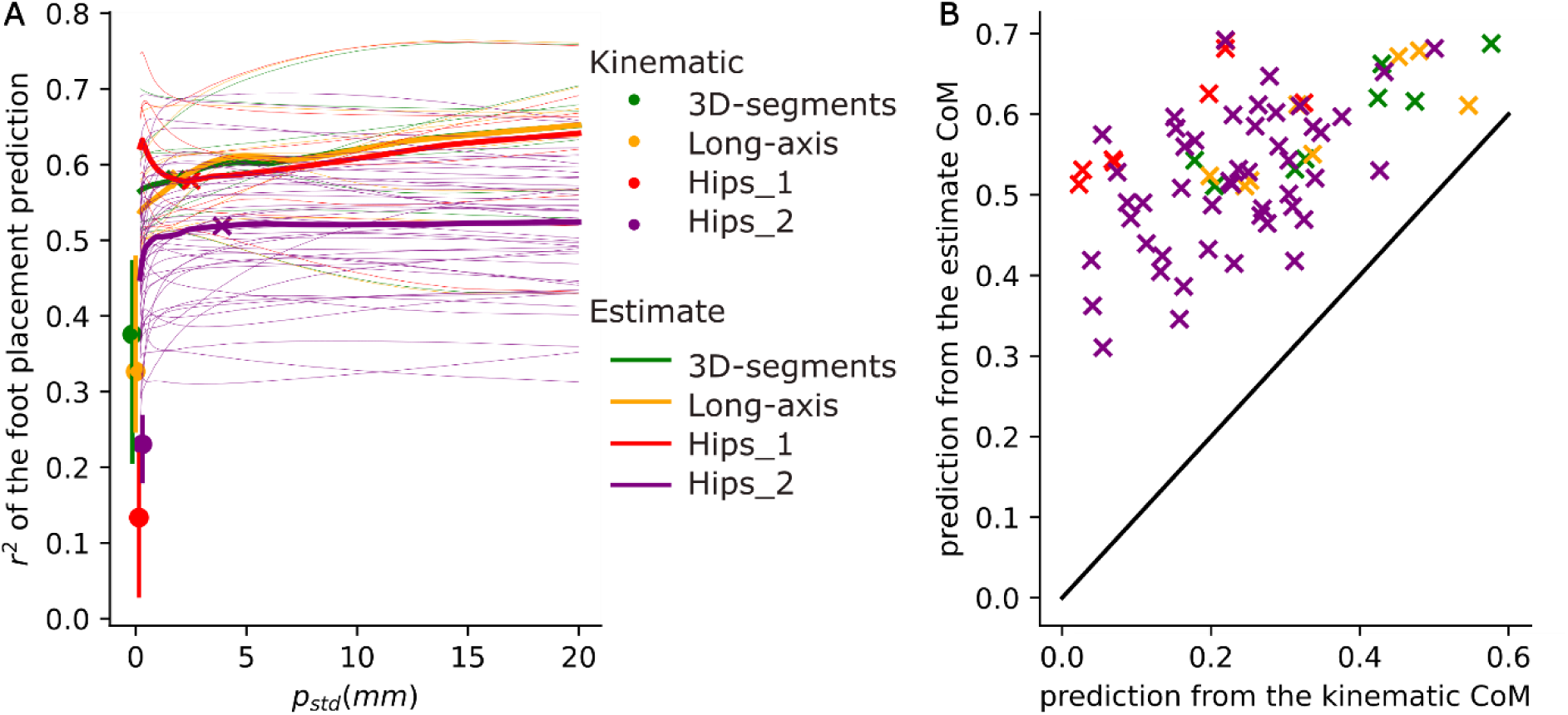
Improved accuracy of the foot placement prediction. r-square of the foot placement prediction for the 3D-segments (green), Long-axis (yellow), Hips_1 (red) and Hips_2 (purple) models. A. The median r-square for the kinematic CoM is indicated as a dot (at *p*_*std*_ = 0) with a bar indicating the 95 % confidence interval. The median r-square for the estimate CoM is indicated as a thick line over the range of *p*_*std*_, with the value at the optimal *p*_*std*_ indicated by a cross. Thin lines correspond to the r-square for individual subjects, speeds and legs. B. r-square of the foot placement prediction from the estimate (at the optimal *p*_*std*_) compared to the kinematic CoM. Each individual subject, speed and leg is indicated by a cross.

**Table 3.** Accuracy of the lateral foot placement prediction. Median and 95 % confidence interval of the r-square of the linear regressions of lateral foot placement onto the CoM state at the preceding flight apex.

|  | 3D-segments | Long-axis | Hips_1 | Hips_2 |
| --- | --- | --- | --- | --- |
| **Kinematics** | 0.38<br>(0.20 to 0.47) | 0.33<br>(0.25 to 0.48) | 0.13<br>(0.03 to 0.22) | 0.23<br>(0.18 to 0.27) |
| **Estimate** | 0.58<br>(0.53 to 0.66) | 0.58<br>(0.52 to 0.68) | 0.58<br>(0.53 to 0.68) | 0.52<br>(0.49 to 0.56) |

## Discussion

### Conclusion

The accuracy of different kinematic models of the whole body center of mass was evaluated by comparing the kinematic CoM acceleration to gravity during the flight phases of running. The advantage of this method is that it does not require pre-processing of the force. This showed a large error (13 to 23 % of gravity for the segmental models). This is of similar magnitude to previous reports of discrepancies between the GRF and the kinematic CoM acceleration of around 10 % of the subject’s weight (Hamner et al., 2010; Futamure et al., 2017; Simonetti et al., 2021; Sturdy et al., 2022). The error was even larger for the hips, reaching 27 to 36 % of gravity. The accuracy of the kinematic models was also evaluated using a classical method: comparing the kinematic position to the kinetic position. As explained in the introduction, this method relies on pre-processing and double integration of the GRF, and therefore cannot be considered as ‘ground truth’. However, it provides a useful estimate of Position Error. We thus found a Position Error Amplitude of 5.8 mm and 6.8 mm for the segmental models, and 12.0 mm and 15.4 mm for the hips. This is of similar magnitude to previous reports of position errors during running ranging from 4.9 mm to 6.8 mm for segmental models and reaching 14.9 mm for the pelvis (Pavei et al., 2017). We confirmed previous findings that the hips provide a very inaccurate measure of the CoM (Whittle, 1997; Eames et al., 1999; Gard et al., 2004; Pavei et al., 2017). Additionally, we found that the accuracy of the 3D-segments model is very close to that of the long-axis model, except for acceleration in the upwards direction (Figure 3). It thus seems that locating the segments’ CoM in the antero-posterior and lateral directions improves the overall CoM accuracy only slightly.

Whereas the error in the kinematic models was high (13 % to 36 % for Acceleration Bias Amplitude and 27 to 56 % for Position Error Amplitude), this error was drastically reduced when using the novel proposed method. Interestingly, the estimate error was very similar for the four kinematic models (1.1 to 3.2 % for both acceleration and position). This suggests that the kinetic signal carries sufficient information to compensate for the imprecision of even the hips model. Thus, while using hips markers alone to determine the CoM position is highly inaccurate, the proposed method combining hip markers and force plate recordings is 4 to 10 times more accurate than the current state-of-the-art kinematic model. The proposed method thus offers a convenient way to have high-accuracy measurements while using few markers.

The accuracy with which the subject’s foot placement could be predicted was critically dependent on the accuracy of the CoM measurement. When using the (inaccurate) hips markers, the CoM state at midstance could only account for 13 to 23 % of the variance in lateral foot placement. When using the more accurate segmental models, this increased to 33 to 38 %. When using the proposed method, this increased to 52 to 58 %. Thus, the poor prediction obtained when using the hips markers does not indicate that the subject has poor control of foot placement, but simply that the CoM measurement is inaccurate.

During walking, balance in the frontal plane is achieved largely through foot placement (for a review, see Bruijn & van Dieën, 2018). The CoM state at midstance can account for 73 % of the variance in lateral foot placement in younger adults (mean age 27.3 years) and only 60 % in healthy older adults (mean age 70.8 years)(Arvin et al., 2018). The r-square of this prediction has therefore been proposed as a measure of balance. When comparing this r-square between two populations, it is crucial that the accuracy of the CoM measurement is the same in the two populations. Indeed, suppose there are no differences in balance control between younger and older subjects, but a single anthropometric table based on measurements of young subjects (de Leva, 1996; Dumas et al., 2007) is used to calculate the CoM for both populations. This will provide a more accurate measure of CoM state in younger subjects, and therefore a better prediction of foot placement, despite there being no actual differences in control. Importantly, the novel method we propose can mitigate biases which are due to the choice of the kinematic model. Indeed, whereas when using only the kinematics, the 3D-segments, Long-axis and Hips_1 models provide values of r-square ranging from 13 % to 38 %, when using the proposed method these three kinematic models provide the same r-square of 58 %. The novel method may therefore be less biased when evaluating balance indicators in populations for which no accurate anthropometric data is available. This improved accuracy is critical for correctly identifying fall risk in pathological populations, and for assessing the progress of rehabilitation.

The method depends on a single parameter: the ratio of position to acceleration noise. Importantly, drastic improvements in accuracy can be obtained without needing to estimate this ratio precisely (Figure 4). In practice the ratio is large. Indeed, good quality force-plates have a measurement noise of a few Newton. With subjects weighing 50 to 100 kg, the force noise is thus on the order of 1 % of the subject’s weight, resulting in an acceleration noise of around 0.01 m/s^2^. The position noise (i.e. the inaccuracy of the kinematic model used to calculate the CoM position) ranges typically from 5 to 15 mm. The sampling frequency of modern motion capture equipment is typically larger than 100 Hz. The squared frequency term results in a large value of *r*. Thus, in practice, the ratio is large and optimal integration relies mainly on double integration of the acceleration at medium and high frequencies (Figure 2.B). Only low frequencies of the position measurement are used (Figure 2.A). This may explain why the accuracy of the estimate is not very sensitive to the kinematic model used. This is qualitatively similar to the heuristic method developed by Maus and colleagues (Maus et al., 2011). However, in our case the cutoff frequency is determined automatically from *r*, and the transfer functions are more complex than a sigmoid (Figure 2, Appendix B section Estimator transfer function).

### Limitations

The two open-access databases used for validating the method only contain ten young subjects. The exact values of the kinematic errors may therefore not match those of the general population. We nevertheless found errors in position which are very similar to previously reported values (Pavei et al., 2017). We only evaluated two of the many segmental models available, and could only use the two subjects of the first dataset to calculate segmental models. We found errors in kinematic position which are similar to previously published results. Importantly, we found that the estimate error was very similar even for models with very different levels of kinematic error. We therefore expect our results to generalize to a wide range of kinematic models.

The Kalman filter yields the optimal combination of position and acceleration measurements, provided that these measurements have additive white Gaussian noise. In practice, the noise structure may be more complex. In particular, force measurements typically have a slow drift. There is no guarantee that the Kalman filter is the optimal way of dealing with such drift. However, our results show that, after removing a linear fit to the force measurement before and after the task, the small residual drift caused an error of less than 1.2 mm in the estimate position (which was much smaller than the error of the double integral of the force, see Supplementary Figure 2).

## Funding

The authors declare that they have received no specific funding for this study.

## Conflict of interest disclosure

The authors declare that they comply with the PCI rule of having no financial conflicts of interest in relation to the content of the article.

## Data, scripts, code, and supplementary information availability

The data was obtained from two publicly available databases. The first database (Wojtusch & von Stryk, 2015) was accessed through the following link:

https://web.sim.informatik.tu-darmstadt.de/humod2/index.html The second database was accessed through the following link: https://doi.org/10.5061/dryad.1nt24m0

The scripts for obtaining the results and figures of this paper are available at: https://github.com/charlotte-lemouel/center_of_mass/tree/main/validation The appendices are available at: https://www.biorxiv.org/content/10.1101/2024.07.24.604923v1.supplementary-material

The code for calculating the kinematic CoM and the optimal combination is available as a python package using: “pip install center_of_mass”. The documentation is available at: https://center-of-mass.readthedocs.io

Alternately, the Python and Matlab code are downloadable at the following link: https://github.com/charlotte-lemouel/center_of_mass

The code for calculating the kinematic CoM is available in Python. It is flexible and works for a variety of marker sets. It requires a minimum of 13 markers - those used in the long-axis model (Tisserand et al., 2016) – to calculate the segments’ CoM location in the longitudinal direction. When additional markers are available, it additionally calculates the segments’ CoM location in the antero-posterior and lateral directions, to provide the best possible accuracy for a given marker set.

The code for calculating the optimal combination is available in both Python and Matlab. The code takes as input the kinetics and the kinematic CoM. The user may use their own method for calculating the kinematic CoM. Based on the results of the current paper, the code uses a default value of *p*_*std*_ = 2 *mm* and *f*_*std*_ = 2 *N*. The user can however manually set the position and force noise. We recommend performing a recording with empty force plates and using the standard deviation of this signal as *f*_*std*_.

## Supporting information

Appendix A - Calculation of the CoM position from kinematics

Appendix B - Center of Mass estimator

## Acknowledgements

Preprint version 3 of this article has been peer-reviewed and recommended by Peer Community in Health and Movement Science (https://doi.org/10.24072/pci.healthmovsci.100153; van Dieen, 2025).

## Supplementary Figures

**Supplementary Figure 1.**
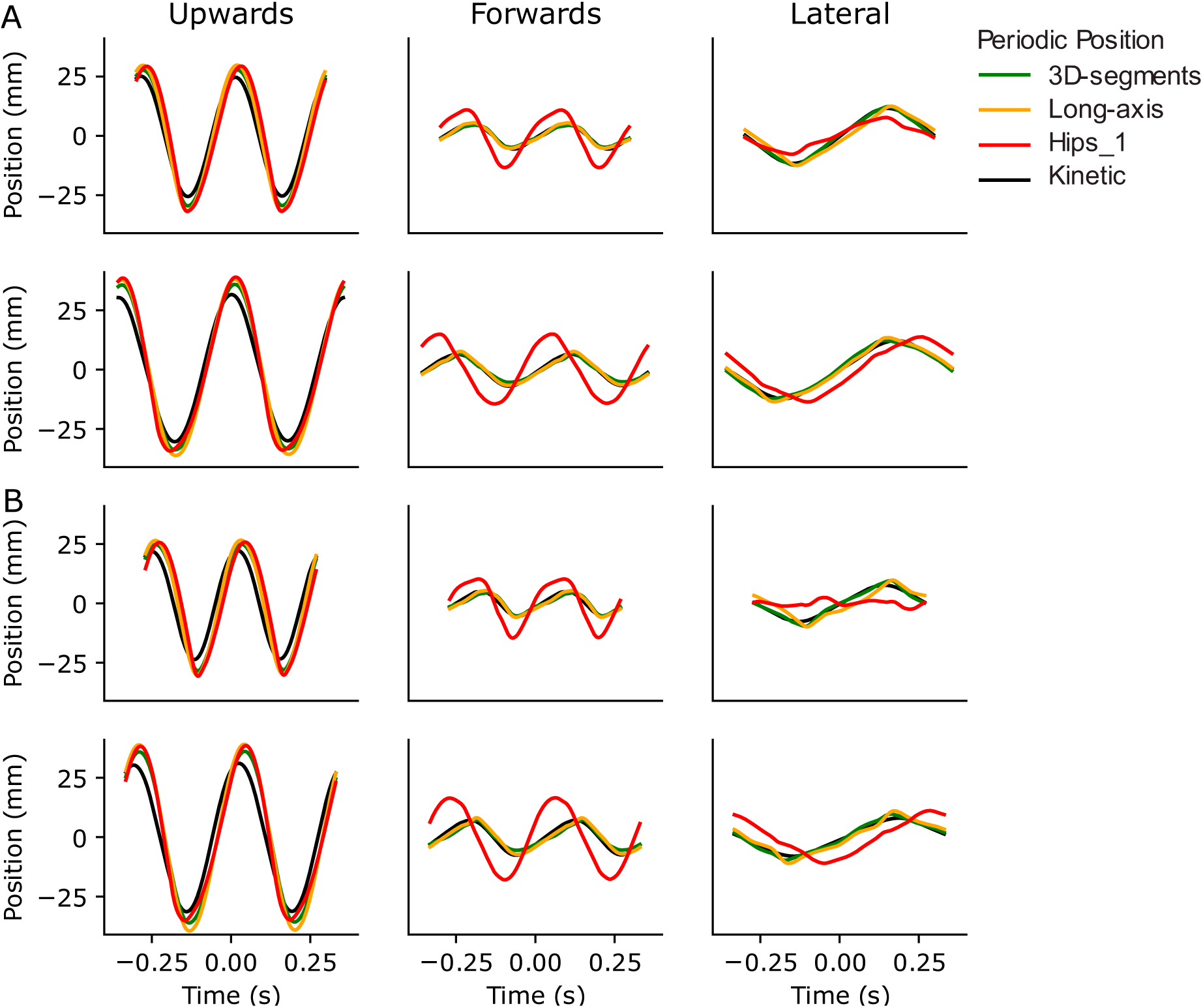
Periodic positions of individual trials. Kinetic periodic position (black) and kinematic periodic positions for the 3D-segments (green), Long-axis (yellow) and Hips_1 (red) models, in the upwards (first column), forwards (middle column) and lateral (right column) directions, when running at 3 m/s (A) and 4 m/s (B) for each of the two subjects, as a function of time relative to heel strike .

**Supplementary Figure 2.**
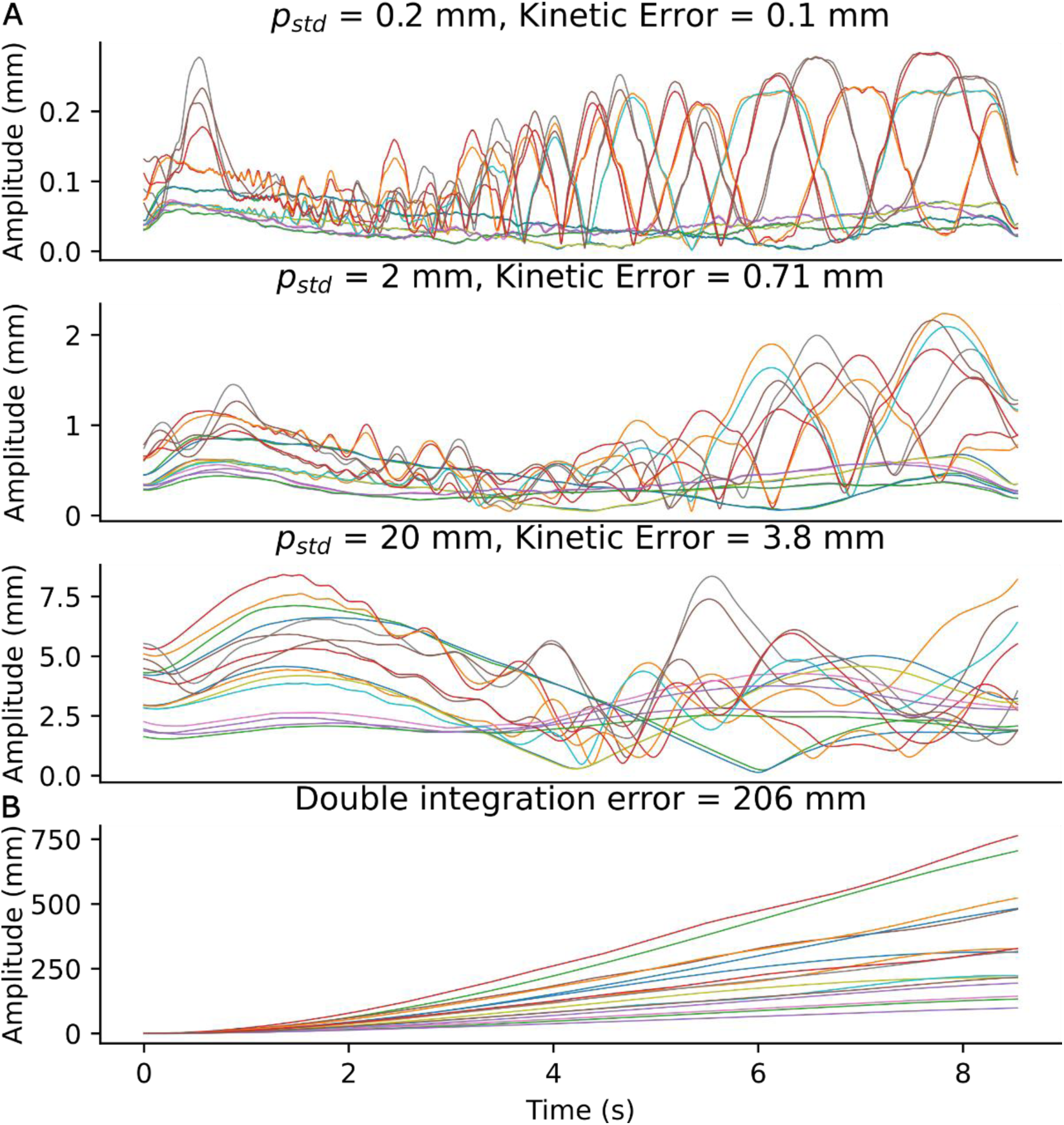
Kinetic Error. Amplitude of the drift in position (A) when the Kalman filter is applied to the empty force plate signals with *p*_*std*_ = 0.2 *mm* (top row), *p*_*std*_ = 2 *mm* (middle row) and *p*_*std*_ = 20 *mm* (bottom row), and (B) after double integration of the empty force plate signal. Individual curves correspond to the initial and final portions of each of the trials of dataset 1. Note that the Kinetic Error of the Kalman filter increases with *p*_*std*_ but does not seem to increase with time, and is much smaller than the drift due to double integration.

## Notes

https://center-of-mass.readthedocs.io

https://github.com/charlotte-lemouel/center_of_mass

### Competing Interest Statement

The authors have declared no competing interest.

### Summary of Updates

The preprint version 3 of this article has been peer-reviewed and recommended by Peer Community in Health and Movement Science (https://doi.org/10.24072/pci.healthmovsci.100153; van Dieen, 2025). The badge of Peer Community in Health and Movement Science has been added to the front page.

