## Appendix A - Calculation of the CoM position from kinematics for "Improved accuracy of the whole body Center of Mass position through Kalman filtering"

Calculation of the human whole body Center of Mass position from kinematic measurements

### I. Overwiew

The calculation of the position of the whole body centre of mass (CoM) is done in two steps.

#### 1. Calculation of the segment coordinate system

First, the body is divided into 16 segments (Figure 1.A, blue ovals), or 15 segments if the thorax and abdomen are merged into a single torso segment. For each segment we calculate the joint centres corresponding to the segment's origin (Figure 1.A, red and yellow dots), the segment coordinate system (SCS) and the segment length $l_s$. We use the conventional SCS recommended by the International Society of Biomechanics (Wu et al., 2002, 2005): when a person stands upright in the anatomical position, with the arms at the sides and the palms facing forwards, the X, Y and Z axes of each segment are oriented forwards, upwards and lateral to the right (Figure 1.B).

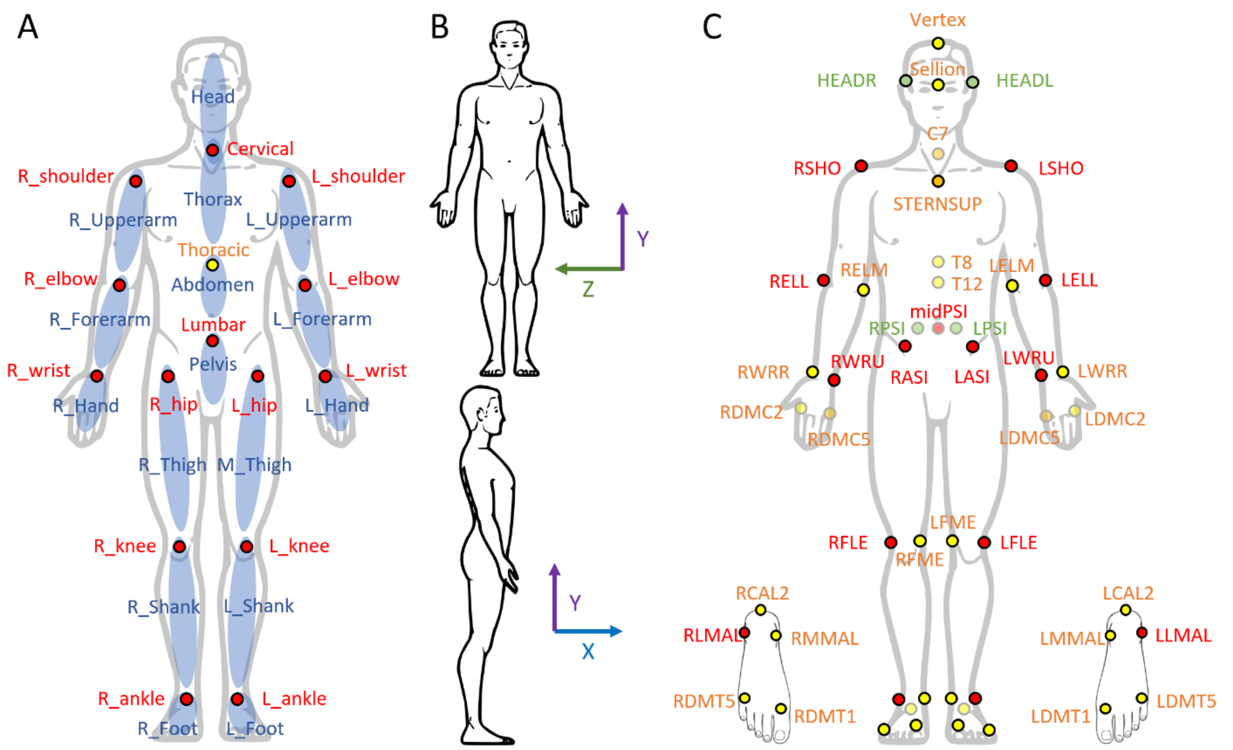

*Figure 1. A. Body segments (blue ovals) and joint centers (red and yellow dots). In the simplified method, the thoracic joint center (yellow) is not available and the abdomen and thorax are merged into a single torso segment. B. In the anatomical posture, the X, Y and Z axes of each segment are oriented forwards, upwards and lateral to the right. C. The simplified method requires 13 markers (red), whereas the reference method uses 37 markers (additional markers in orange and yellow). Alternative markers (green) can be used to calculate the head and pelvis SCS.*

To calculate the joint centers and SCS, the reference method (Dumas et al., 2007a, 2007b, 2015) uses 37 anatomical markers (Figure 1.C, markers in red, orange and yellow). A simplified method was proposed (Tisserand et al., 2016) which requires only 13 markers (Figure 1.C, markers in red). The code is written in a modular fashion, such that when the complete set of markers is available for a segment, the reference method is applied to that segment. Otherwise, we use either the simplified method or an intermediate version using the markers in orange (Figure 1.C), described when applicable. In addition, a method is provided to calculate the pelvis and head SCS from alternative markers (Figure 1.C, markers in green). The marker names and definitions are in Table 1.

When the length $l_{distal}$ of a distal segment (head, hand or foot) cannot be estimated from the markers present, it is estimated from the length $l_{proximal}$ of the segment proximal to it, according to (Tisserand et al., 2016):

$$l_{distal} = l_{proximal} \frac{l_{distal}^{ref}}{l_{proximal}^{ref}}$$

The reference lengths are taken from (Dumas et al., 2007a) Table 2 for the torso, from (Dumas and Wojtusch, 2018) for the other segments, and reported in Table 2.

*Table 1 **Marker names and definitions**. The markers used in the simplified method are in red. Additional markers for the intermediate and references methods are in orange and yellow. Alternative markers are in green*

| Segment | Marker name | Marker definition |
|---|---|---|
| Pelvis | LASI / RASI | Left / Right anterior iliac spines |
| | LPSI / RPSI | Left / Right posterior iliac spines |
| | midPSI | midpoint between the left and right posterior iliac spines |
| Trunk | T8 | 8th thoracic vertebra |
| | T12 | 12th thoracic vertebra |
| | C7 | 7th cervical vertebra |
| | STERNSUP | suprasternal notch (on the upper end of the sternum, below the two ends of the claviculae) |
| | LSHO, RSHO | Left / Right acromion (bony prominence) |
| Head and neck | Vertex | Head vertex |
| | Sellion | Head sellion |
| | HEADL / HEADR | Either: Left / Right tragion<br>Or: markers placed symmetrically Left / Right on a headband |
| Upper arm | LELL / RELL | Left / Right lateral humeral epicondyle |
| | LELM / RELM | Left / Right medial humeral epicondyle |
| Forearm | LWRU / RWRU | Left / Right ulnar styloid process |
| | LWRR / RWRR | Left / Right radial styloid process |
| Hand | LDMC5 / RDMC5 | Left / Right distal metacarpal V (on the base of the finger) |
| | LDMC2 / RDMC2 | Left / Right distal metacarpal II (on the base of the finger) |
| Thigh | LFLE / RFLE | Left / Right lateral femoral epicondyle |
| | LFME / RFME | Left / Right medial femoral epicondyle |
| Shank | LLMAL / RLMAL | Left / Right lateral malleolus |
| | LMMAL / RMMAL | Left / Right medial malleolus |
| Foot | LDMT1 / RDMT1 | Left / Right distal metatarsal I |
| | LDMT5 / RDMT5 | Left / Right distal metatarsal V |
| | LCAL2 / RCAL2 | Left / Right tuber calcanei (most prominent point of th heel) |

2. Regression equations

Then, for each segment, the segment origin $O_s$, SCS $(X_s, Y_s, Z_s)$ and length $l_s$ are used to calculate the position of the segment's CoM according to the following regression equation:

$$CoM_s = O_s + \frac{l_s}{100}(x_s X_s + y_s Y_s + z_s Z_s)$$

The length percents $x_s, y_s, z_s$ are provided in Table 2.
Finally, the whole body CoM is calculated as the barycenter of the different segments, weighted by the percent of the segments' mass relative to the total body mass $m_s$ (Table 2), according to the following regression equation:

$$CoM_{tot} = \sum_s m_s CoM_s$$

The length percents $x_s, y_s, z_s$ and mass percents $m_s$ are obtained according to the method of (Dumas et al., 2007a, 2007b, 2015). This method uses photogrammetry data of 31 males (mean age 27.5 years old, mean weight 77.3 kg, mean stature 1.77 m) (McConville et al., 1980) and 46 females (mean age 31.2 years old, mean weight 63.9 kg, mean stature 1.61 m) (Young et al., 1983). The values are summarised in (Dumas and Wojtusch, 2018), except for the values for the torso segment, which are taken from (Dumas et al., 2007a) Table 2.

*Table 2 Anthropometric values used for the regression equations (Dumas et al. 2007a, 2007b, 2015)*

| Segment | Reference | Sex | Position of the segment CoM (% of the segment length) | | | Segment mass (% of the total body mass) | Segment length (mm) |
|---|---|---|---|---|---|---|---|
| | | | Forwards (X) | Upwards (Y) | Rightwards (Z) | | |
| Head and neck | Dumas & Wojtusch (2018), Table 1 | female | 0.8 | 55.9 | -0.1 | 6.7 | 243 |
| | | male | 2 | 53.4 | 0.1 | 6.7 | 278 |
| Thorax | Dumas & Wojtusch (2018), Table 2 | female | 1.5 | -54.2 | 0.1 | 26.3 | 322 |
| | | male | 0 | -55.5 | -0.4 | 30.4 | 334 |
| Abdomen | Dumas & Wojtusch (2018), Table 3 | female | 21.9 | -41 | 0.3 | 4.1 | Not used in the code |
| | | male | 17.6 | -36.1 | -3.3 | 2.9 | |
| Torso | Dumas et al. (2007a), Table 2 | female | -1.6 | -43.6 | -0.6 | 30.4 | 429 |
| | | male | -3.6 | -42 | -0.2 | 33.3 | 477 |
| Pelvis | Dumas & Wojtusch (2018), Table 4 | female | -7.2 | -22.8 | 0.2 | 14.7 | Not used in the code |
| | | male | -0.2 | -28.2 | -0.6 | 14.2 | |
| Upper arm | Dumas & Wojtusch (2018), Table 5 | female | -5.5 | -50 | -3.3 | 2.3 | |
| | | male | 1.8 | -48.2 | -3.1 | 2.4 | |
| Forearm | Dumas & Wojtusch (2018), Table 6 | female | 2.1 | -41.1 | 1.9 | 1.4 | 247 |
| | | male | -1.3 | -41.7 | 1.1 | 1.7 | 283 |
| Hand | Dumas & Wojtusch (2018), Table 7 | female | 7.7 | -76.8 | 4.8 | 0.5 | 71 |
| | | male | 8.2 | -83.9 | 7.5 | 0.6 | 80 |
| Thigh | Dumas & Wojtusch (2018), Table 8 | female | -7.7 | -37.7 | 0.8 | 14.6 | 388 |
| | | male | -4.1 | -42.9 | 3.3 | 12.3 | 433 |
| Shank | Dumas & Wojtusch (2018), Table 9 | female | -4.9 | -40.4 | 3.1 | 4.5 | 117 |
| | | male | -4.8 | -41 | 0.7 | 4.8 | 139 |

#### II. Trunk and head segments

##### 1. Pelvis: hip and lumbar joint centers

The markers required for the calculation of the hip and lumbar joint centers (Figure 2) are: the left and right anterior iliac spines (LASI, RASI), and the midpoint between the left and right posterior iliac spines (midPSI).

**Pelvis coordinate system** (Wu et al., 2002)
The Z direction is parallel to the line from LASI to RASI, pointing rightwards.
The Y direction is orthogonal to the plane containing LASI, RASI and midPSI, pointing upwards.
The X direction is orthogonal to the Y and Z directions, pointing forwards.

**Hip and lumbar joint centers**
Based on measurements of the pelvic bones of 28 “small female” and 33 “midsize male” subjects aged 18 to 55 years old (Reynolds et al., 1982), a method was developed to calculate the hip and lumbar joint centers from anatomical markers placed on the subject’s skin (Reed et al., 1999). This method was then adapted (Dumas and Wojtusch, 2018) to provide the position of these joint centers in the ISB pelvic reference frame. These positions are given relative to the midpoint between LASI and RASI. The distances are expressed as a percent of the pelvis width (% PW), with pelvis width defined as the distance between LASI and RASI. The pelvis width is calculated as the median of the distance between LASI and RASI throughout the trial.

*Table 3 **Pelvis**. Positions of the lumbar and hip joint centers (% pelvis width – PW), taken from Dumas and Wojtusch (2018), Table 3 for the lumbar joint center, and Table 4 for the hip joint centers.*

| | | Forwards (X) | Upwards (Y) | Rightwards (Z) |
|---|---|---|---|---|
| Lumbar joint center | Female | - 34.0 % PW | 4.9 % PW | 0 % PW |
| | Male | - 33.5 % PW | - 3.2 % PW | 0 % PW |
| Right / Left hip joint center | Female | - 13.9 % PW | - 33.6 % PW | +/- 37.2 % PW |
| | Male | - 9.5 % PW | - 37.0 % PW | +/- 36.1 % PW |

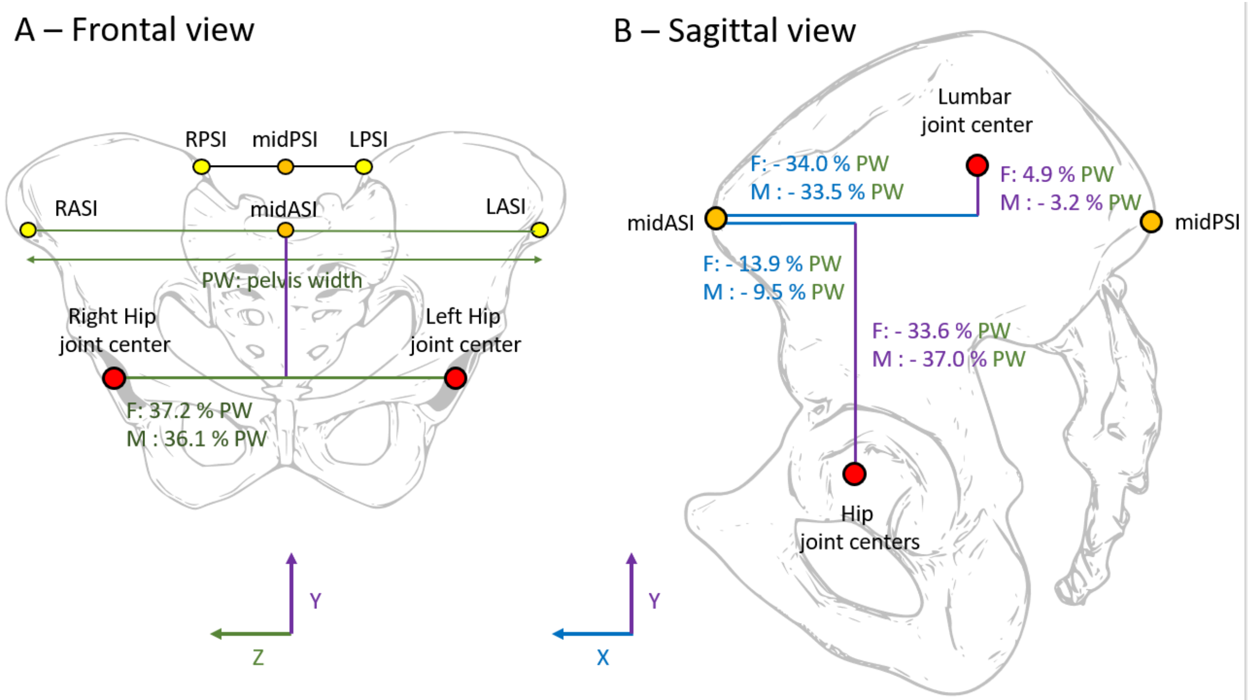

*Figure 2. **Pelvis**. Positions of the lumbar and hip joint centers (red) relative to the anatomical markers (yellow) in the frontal (A) and sagittal (B) planes.*

**Pelvis SCS origin and length**

The segment origin is the lumbar joint center, and the segment length is the distance between the lumbar joint center and the projection of the hip joint centers in the pelvic sagittal plane (Dumas et al., 2007a), Table 2.

2. Torso

Depending on the markers present, the torso is either decomposed into two segments (thorax and abdomen) or considered as a single segment.

a. Thorax and abdomen

If the 8th and 12th thoracic vertebra (T8 and T12), the 7th cervical vertebra (C7) and the suprasternal notch (STERNSUP) are present, then the trunk is decomposed into two segments : abdomen and thorax (Dumas et al., 2015; Dumas and Wojtusch, 2018).

The thoracic joint center (Figure 3, left) is calculated according to the method of (Reed et al., 1999) for males, adapted for females by (Dumas et al., 2015) using the data from (Robbins, 1983). The thoracic sagittal plane is defined as the plane containing T8, C7 and STERNSUP (Dumas and Wojtusch, 2018), Table 2). The thorax width is defined as the distance between C7 and STERNSUP. The thoracic joint center is in the thoracic sagittal plane, forwards of T12 at a distance of 50 % of thorax width for females (52 % for males), and at an angle of 92 ° for females (94 ° for males) from the line joining T12 and T8 (Dumas and Wojtusch, 2018), Table 2).

The cervical and shoulder joint centers (Figure 3, right) are calculated according to (Dumas et al., 2007a), using the method of (Reed et al., 1999) and the data of (Robbins, 1983) from 25 midsize males and 25 females. The cervical joint center is located in the thorax sagittal plane, forwards and upwards of C7 by a distance of 53 % of the thorax width for females (55 % for males) and at an angle of 14° for females (8° for males) relative to the line joining C7 and STERNSUP.

The right (or left) shoulder joint center is located in the plane parallel to the thoracic sagittal plane and containing the right (or left) acromion. It is forwards and downwards of the acromion by 36 % of thorax width for females (33 % for males) and at an angle of 5° for females (11° for males) to the line joining C7 and STERNSUP.

**Thorax coordinate system**
The Z direction is orthogonal to the plane containing T8, C7 and STERNSUP, pointing rightwards.
The Y direction is parallel to the line from the thoracic joint center to the cervical joint center, pointing upwards.
The X direction is orthogonal to the Y and Z directions, pointing forwards.
The origin is at the cervical joint center.
The length is the distance from the cervical to the thoracic joint center.

**Abdomen coordinate system**
The Y direction is parallel to the line from the lumbar joint center to the thoracic joint center, pointing upwards.
The X direction is the average between the X directions of the pelvis and thorax, projected onto the plane orthogonal to the Y direction of the abdomen and normalised.
The Z direction is orthogonal to the X and Y directions, pointing rightwards.
The origin is at the thoracic joint center.
The length is the distance from the thoracic to the lumbar joint center.

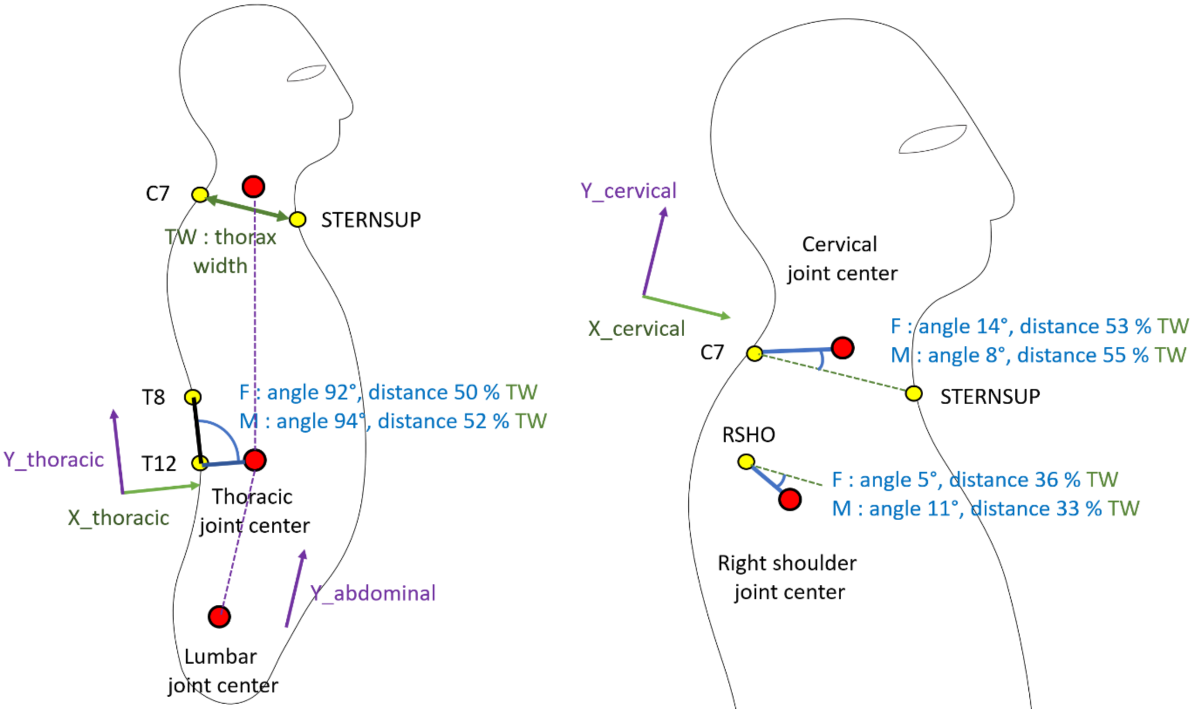

*Figure 3. **Torso**. Positions of the thoracic, cervical and shoulder joint centers*

b. Torso segment

If T8 and T12 are not both present, then the position of the thoracic joint center cannot be calculated, and the abdomen and thorax are merged into a single segment, the torso, as in (Dumas et al., 2007a).
If C7 and STERNSUP are both present, the cervical and shoulder joint centers are calculated as described previously with one modification: the sagittal plane is defined as the plane containing C7, STERNSUP and the lumbar joint center.

**Torso coordinate system**
The Y direction is parallel to the line from the lumbar joint center to the cervical joint center, pointing upwards.
The Z direction is orthogonal to the plane containing C7, STERNSUP and the lumbar joint center, pointing rightwards.
The X direction is orthogonal to the Y and Z directions, pointing forwards.
The origin is at the cervical joint center.
The length is the distance from the cervical to the lumbar joint center.

c. Simplified torso segment

If STERNSUP is not present, the shoulder joint centers are located at the acromion markers. If C7 is present, the cervical joint center is located at C7, as proposed by (de Leva, 1996). Otherwise, it is at the midpoint between the shoulder joint centers, as proposed by (Tisserand et al., 2016).

**Simplified torso coordinate system**
Only the Y direction is available. It is parallel to the line from the lumbar joint center to the cervical joint center, pointing upwards.
The origin is at the cervical joint center.
The length is the distance from the cervical to the lumbar joint center.

3. Head and neck segment

The segment origin is the cervical joint center.

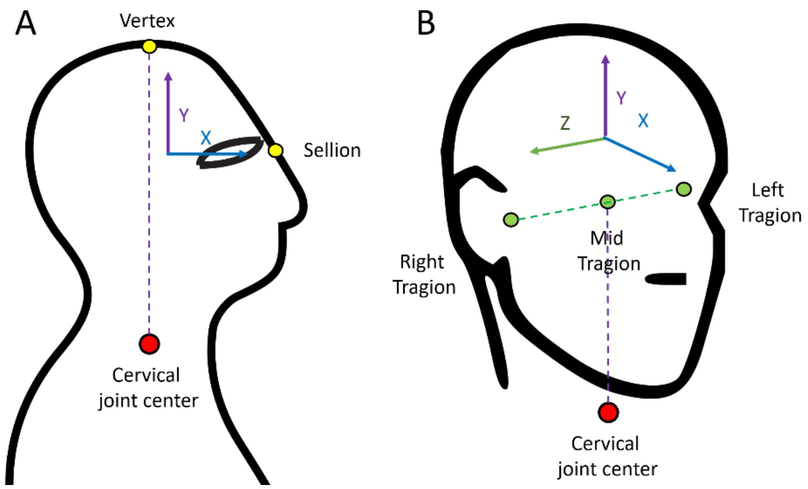

*Figure 4. **Head.** SCS in the reference method (A) and using alternative markers (B).*

a. Reference head and neck SCS

When the head vertex and sellion markers are present (Figure 4.A), the head coordinate system is defined as in (Dumas and Wojtusch, 2018):
The Y direction is parallel to the line from the cervical joint center to the vertex, pointing upwards.
The Z direction is orthogonal to the plane containing the cervical joint center, the vertex and the sellion, pointing rightwards.
The X direction is orthogonal to the Y and Z directions, pointing forwards.
The segment length is the distance between the cervical joint center and the vertex.

b. Alternative head and neck SCS

When, instead of the sellion and vertex, either the left and right tragion are provided, or a headband is used which has two symmetrically placed markers on the left and right (Head_L and Head_R), the head coordinate system is defined as follows (Figure 4.B):
The Y direction is parallel to the line from the cervical joint center to the midpoint between Head_L and Head_R, pointing upwards.
The X direction is orthogonal to the plane containing the cervical joint center and Head_L and Head_R, pointing forwards.
The Z direction is orthogonal to X and Y directions, pointing rightwards.
The segment length is scaled by the length of the proximal segment (either the thorax if it is available, or the torso otherwise):

$$l_{head} = l_{thorax/torso} \frac{l_{head}^{ref}}{l_{thorax/torso}^{ref}}$$

c. Simplified head and neck SCS

If head markers are not present, similarly to (Tisserand et al., 2016), the head (X,Y,Z)-axes are assumed to match those of either the thorax (if it is available) or torso (otherwise). Note that in the simplified method originally proposed by (Tisserand et al., 2016), only the Y direction is available for the torso, therefore only the Y direction is available for the head.
The segment length is scaled by the length of the proximal segment (either the thorax if it is available, or the torso otherwise):

$$l_{head} = l_{thorax/torso} \frac{l_{head}^{ref}}{l_{thorax/torso}^{ref}}$$

### III. Limbs

#### 1. Overview

The coordinate systems of the limb segments are all calculated in a similar manner (except for the feet). The segment origin is the proximal joint center and the segment length is the distance from the proximal to the distal joint center.

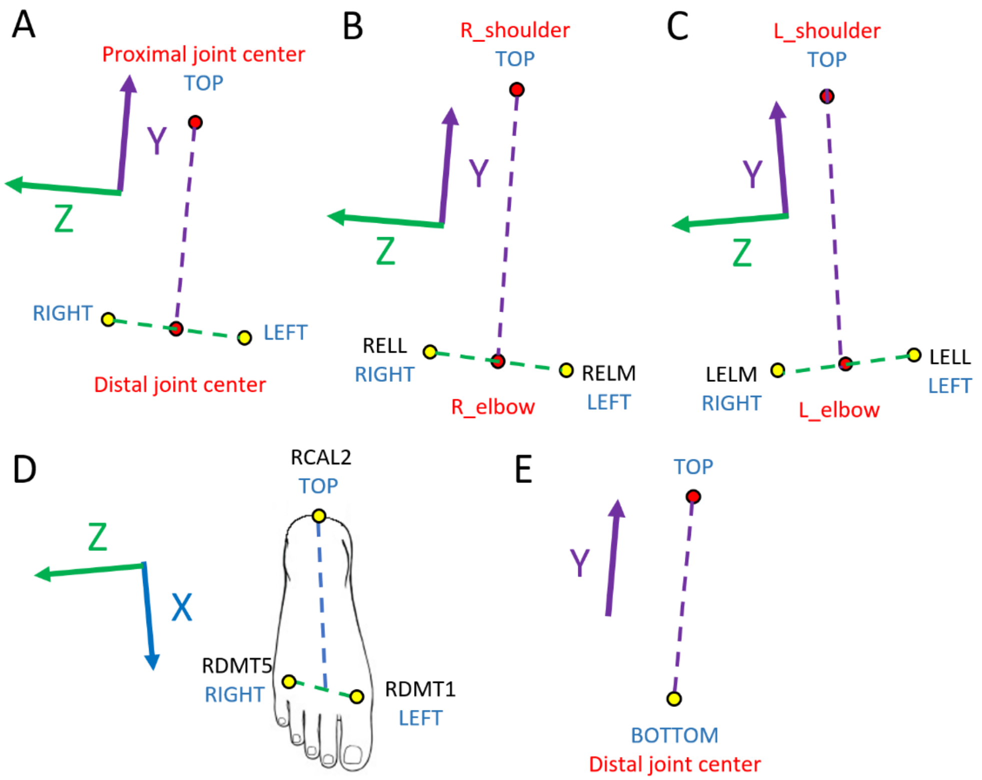

*Figure 5. **Limb segment coordinate systems**. In the reference method (A-D), the SCS is calculated from the proximal joint center (TOP) and two distally placed markers (RIGHT and LEFT). For right limb segments (B) versus left limb segments (C), the lateral and medial markers are flipped. D. SCS of the right foot. E. SCS in the simplified method.*

a. Reference method

The reference method uses the proximal joint center (TOP, Figure 5.A-C) and two distal markers placed medially and laterally of the distal joint center (RIGHT and LEFT, Figure 5.A-C). The distal joint center is placed at the mid-point between the two distal markers (RIGHT and LEFT). The Y axis is from the distal to the proximal joint center. The X axis is orthogonal to the plane containing the proximal joint center and the two distal markers, pointing forwards. The Z axis is orthogonal to X and Y, pointing rightwards.

For example, for the upper arm, the proximal joint is the shoulder joint center (TOP, Figure 5.B-C). For the right upper arm (Figure 5.B), the rightwards distal marker (RIGHT) is the right humeral lateral epicondyle (RELL) and the leftwards one is the right humeral medial epicondyle (RELM). For the left upper arm, the lateral and medial distal markers are flipped (Figure 5.C): RIGHT corresponds to the left humeral lateral epicondyle (LELM) and the LEFT to the left humeral medial epicondyle (LELL). In this way, when the person stands in the reference anatomical posture, the Z-axis of all limb segments is oriented rightwards.

Note that for the feet (Figure 5.D), the calcaneus marker on the heel (LCAL2 and RCAL2) is taken as the TOP marker rather than the ankle joint (Dumas and Wojtusch 2018, Table 10). The X axis (forwards) - rather than the Y axis (upwards) - is from the ankle to the toe joint.

b. Simplified methods

If a single distal marker is available (Figure 5.E, BOTTOM), we apply a simplified method (Tisserand et al. 2016): the distal joint center is assumed to be at the BOTTOM marker. Only the Y-axis is available, from the distal to the proximal joint center.

For the hands (respectively feet), if no distal marker is present, the segment coordinate system is assumed to match that of the forearm (respectively shank), and the segment length is scaled by the length of the proximal segment.

2. Upper arm

**Reference SCS**
The elbow joint center is the midpoint between the medial and lateral humeral epicondyles (ELM and ELL).
The Y direction is parallel to the line from the elbow to the shoulder joint center, pointing upwards.
The Z direction is orthogonal to Y, in the frontal plane containing the shoulder joint center, ELM and ELL, pointing rightwards.
The X direction is orthogonal to Y and Z, pointing forwards.

**Simplified SCS**
If only ELL is available, the elbow joint center is at ELL, and only the Y direction is available, defined as above.

3. Forearm

**Reference SCS**
The wrist joint center is the midpoint between the radial and ulnar styloids (WRR and WRU).
The Y direction is parallel to the line from the wrist to the elbow joint center, pointing upwards.
The Z direction is orthogonal to Y, in the frontal plane containing the elbow joint center, WRR and WRU, pointing rightwards.
The X direction is orthogonal to Y and Z, pointing forwards.

**Simplified SCS**
If only WRU is available, the wrist joint center is at WRU, and only the Y direction is available, defined as above.

4. Hand

**Reference SCS**
The finger joint center is the midpoint between the 2nd and 5$^{th}$ metacarpal heads (DMC2 and DMC5).
The Y direction is parallel to the line from the finger to the wrist joint center, pointing upwards.
The Z direction is orthogonal to Y, in the frontal plane containing the wrist joint center, DMC2 and DMC5, pointing rightwards.
The X direction is orthogonal to Y and Z, pointing forwards.

**Simplified SCS**
If hand markers are not available, the hand (X,Y,Z)-axes are assumed to match those of the forearm. Note that in the simplified method proposed by Tisserand et al. (2016), only the Y direction of the forearm is available.
The segment length is scaled by the length of the forearm:

$$l_{hand} = l_{forearm} \frac{l_{hand}^{ref}}{l_{forearm}^{ref}}$$

5. Thigh

**Reference SCS**
The knee joint center is the midpoint between the lateral and medial femoral epicondyles (FLE and FME).
The Y direction is parallel to the line from the knee to the hip joint center, pointing upwards.
The Z direction is orthogonal to Y, in the frontal plane containing the hip joint center, FLE and FME, pointing rightwards.

The X direction is orthogonal to Y and Z, pointing forwards.

**Simplified SCS**
If only FLE is available, the knee joint center is at FLE, and only the Y direction is available, defined as above.

6. Shank

**Reference SCS**
The ankle joint center is the midpoint between the lateral and medial femoral malleolus (LMAL and MMAL).
The Y direction is parallel to the line from the ankle to the knee joint center, pointing upwards.
The Z direction is orthogonal to Y, in the frontal plane containing the knee joint center, LMAL and MMAL, pointing rightwards.
The X direction is orthogonal to Y and Z, pointing forwards.

**Simplified SCS**
If only LMAL is available, the ankle joint center is at LMAL, and only the Y direction is available, defined as above.

7. Foot

**Reference SCS**
The toe joint center is the midpoint between the 1st and 5th metatarsal heads (DMT1 and DMT5).
The X direction is parallel to the line from the calcaneus to the toe joint center, pointing forwards.
The Z direction is orthogonal to X, in the transverse plane containing the calcaneus, DMT1 and DMT5, pointing rightwards.
The Y direction is orthogonal to X and Z, pointing upwards.
The origin is the ankle joint center and the segment length is the distance from the ankle to the toe joint center (Dumas et al., 2007a).

**Simplified SCS**
If feet markers are not available, then the feet (X,Y,Z)-axes are assumed to match those of the lower leg.
The segment length is scaled by the length of the shank:

$$l_{foot} = l_{shank}\,\frac{l_{foot}^{ref}}{l_{shank}^{ref}}$$
