## Appendix B - Center of Mass estimator for "Improved accuracy of the whole body Center of Mass position through Kalman filtering"

### Contents

**State dynamics** 1

**Measurements** 2

**State estimator** 3

**Optimal estimator gain** 3

**Steady state** 5

### State dynamics

Each dimension is considered independently. The position $x$ and acceleration $a$ are related according to :

$$\frac{d^2}{dt^2}x = a$$

We introduce the speed $v$, and rewrite this as a first order linear differential equation on the system state $X = \begin{pmatrix} x \\ v \end{pmatrix}$

$$\frac{d}{dt}x = v$$

$$\frac{d}{dt}v = a$$

$$\frac{d}{dt}\begin{pmatrix} x \\ v \end{pmatrix} = \begin{pmatrix} 0 & 1 \\ 0 & 0 \end{pmatrix}\begin{pmatrix} x \\ v \end{pmatrix}(t) + \begin{pmatrix} 0 \\ 1 \end{pmatrix}a(t)$$

The observations of the state are made at discrete measurement timepoints separated by a duration T = 1/F, where F is the sampling frequency. To obtain the change in state between successive measurements, we integrate the previous equation, assuming that the acceleration $a$ is constant between measurements:

$$\begin{pmatrix} x \\ v \end{pmatrix} (kT + T) = \begin{pmatrix} 1 & T \\ 0 & 1 \end{pmatrix} \begin{pmatrix} x \\ v \end{pmatrix} (kT) + \begin{pmatrix} \frac{T^2}{2} \\ T \end{pmatrix} a\,(kT)$$

We introduce the notations: $x_k = x\,(kT)$ , $v_k = v\,(kT)$ , $a_k = a\,(kT)$, $X_k = \begin{pmatrix} x_k \\ v_k \end{pmatrix}$.

The discrete time dynamics are then:

$$x_{k+1} = x_k + Tv_k + \frac{T^2}{2} a_k$$

$$v_{k+1} = v_k + Ta_k$$

$$X_{k+1} = \begin{pmatrix} x_{k+1} \\ v_{k+1} \end{pmatrix} = \begin{pmatrix} 1 & T \\ 0 & 1 \end{pmatrix} X_k + \begin{pmatrix} \frac{T^2}{2} \\ T \end{pmatrix} a_k$$

We introduce the matrix $A = \begin{pmatrix} 1 & T \\ 0 & 1 \end{pmatrix}$ and the vector $B = \begin{pmatrix} \frac{T^2}{2} \\ T \end{pmatrix}$. The dynamics are then given by:

$$X_{k+1} = AX_k + Ba_k \tag{1}$$

### Measurements

The position measurement $x_k^{meas}$ is given by:

$$x_k^{meas} = x_k + w_k = CX_k + w_k \tag{2}$$

where $C = \begin{pmatrix} 1 & 0 \end{pmatrix}$ and the position noise $w_k$ is assumed to be Gaussian with zero mean and standard deviation $p_{std}$.

The acceleration measurement $a_k^{meas}$ is given by:

$$a_k^{meas} = a_k + \nu_k$$

where the acceleration noise $\nu_k$ is assumed to be Gaussian with zero mean and standard deviation $a_{std}$.

#### State estimator

We define $X_k^{est} = \begin{pmatrix} x_k^{est} \\ v_k^{est} \end{pmatrix}$ as the best estimate of the system state $X_k$ given all the observations $a_{j=1,..,k}^{meas}, x_{j=1,..,k}^{meas}$up to timepoint k. Given this estimate $X_k^{est}$ and the acceleration measurement $a_k^{meas}$, a prediction $X_{k+1}^{pred}$ of the system state at timepoint k+1 can be obtained by double integration of the measured acceleration (as in equation 1) :

$$X_{k+1}^{pred} = AX_k^{est} \ + Ba_k^{meas} \tag{3}$$

The position prediction $x_{k+1}^{pred}$ is then compared to the position measurement $x_{k+1}^{meas}$, and the position error $x_{k+1}^{meas} - x_{k+1}^{pred}$ is used to adjust the estimate of both position and speed at timepoint k+1 using the estimator gain $L_k$:

$$X_{k+1}^{est} = X_{k+1}^{pred} + L_k \left( x_{k+1}^{meas} - x_{k+1}^{pred} \right) = X_{k+1}^{pred} + L_k x_{k+1}^{meas} - L_k C X_{k+1}^{pred} = (1 - L_k C) X_{k+1}^{pred} + L_k x_{k+1}^{meas}$$

Replacing equations 3 and **??** in the above equation, we obtain the following dynamics for the state estimator:

$$X_{k+1}^{est} = (1 - L_k C)(AX_k^{est} + Ba_k^{meas}) + L_k(CX_{k+1} + w_{k+1}) \tag{4}$$

#### Optimal estimator gain

We introduce the estimation error at timestep k: $\Delta_k = X_k - X_k^{est}$, and the mean squared error $P_k = \mathrm{E}\left(\Delta_k . \Delta_k^T\right)$, where E(.) denotes the expected value. We wish to find the estimator gain $L_k$ which minimises $P_{k+1}$.

First, we determine the evolution of the estimation error using equation **??**:

$$\begin{aligned} \Delta_{k+1} &= X_{k+1} - X_{k+1}^{est} \\ &= X_{k+1} - (1 - L_k C)\left(AX_k^{est} + Ba_k^{meas}\right) - L_k\left(CX_{k+1} + w_{k+1}\right) \\ &= (1 - L_k C)\left(X_{k+1} - AX_k^{est} - Ba_k^{meas}\right) - L_k w_{k+1} \end{aligned}$$

Replacing equation 1 in the above equation, we obtain:

$$\begin{aligned} \Delta_{k+1} &= (1 - L_k C)\left(AX_k + Ba_k - AX_k^{est} - Ba_k^{meas}\right) - L_k w_{k+1} \\ &= (1 - L_k C)\left(A(X_k - X_k^{est}) + B(a_k - a_k^{meas})\right) - L_k w_{k+1} \\ &= (1 - L_k C)\left(A\Delta_k - B\nu_k\right) - L_k w_{k+1} \end{aligned}$$

To obtain the evolution of the mean squared error $P_{k+1} = \mathrm{E}\left(\Delta_{k+1}.\Delta_{k+1}^T\right)$, we use the fact that the measurement errors $\nu_k$ and $w_k$ are independent from each other and independent of the current estimation error:

$$0 = \mathrm{E}\left(\Delta_k.\nu_k^T\right) = \mathrm{E}\left(\Delta_k.w_k^T\right) = \mathrm{E}\left(w_k.\nu_k^T\right)$$

We then have

$$\begin{aligned}
P_{k+1} &= (1 - L_k C) A \mathrm{E}\left(\Delta_k.\Delta_k^T\right) A^T \left(1 - C^T L_k^T\right) \\
&+ (1 - L_k C) B \mathrm{E}\left(\nu_k.\nu_k^T\right) B^T \left(1 - C^T L_k^T\right) \\
&+ L_k \mathrm{E}\left(w_k.w_k^T\right) L_k^T \\
&= (1 - L_k C) A P_k A^T \left(1 - C^T L_k^T\right) + a_{std}^2 (1 - L_k C) B B^T \left(1 - C^T L_k^T\right) + p_{std}^2 L_k L_k^T
\end{aligned}$$

To find $L_k$ which minimises $P_{k+1}$, we rewrite the above equation as a second order polynomial in $L_k$:

$$P_{k+1} = A P_k A^T + a_{std}^2\ B B^T + L_k \left(-C A P_k A^T - a_{std}^2 C B B^T\right)$$
$$+ \left(-A P_k A^T C^T - a_{std}^2 B B^T C^T\right) L_k^T + L_k (C A P_k A^T C^T + a_{std}^2 C B B^T C^T + p_{std}^2) L_k^T \quad (5)$$

To simplify notations, we introduce the symmetrical matrix $M_k$ corresponding to the mean squared error of the predictor:

$$M_k = A P_k A^T + a_{std}^2 B B^T = E\left((X_k - X_k^{pred}).\left(X_k - X_k^{pred}\right)^T\right) \quad (6)$$

Replacing in the equation 26, we obtain:

$$P_{k+1} = M_k - L_k C M_k - M_k C^T L_k^T + L_k (C M_k C^T + p_{std}^2) L_k^T \quad (7)$$

To determine the value of $L_k$ which minimises equation 7, we introduce the scalar $r_k$ corresponding to the sum of the variances of the position measurement and position prediction:

$$r_k = C M_k C^T + p_{std}^2$$

Replacing in equation 7, we obtain:

$$P_{k+1} = M_k + r_k \left(-L_k \frac{C M_k}{r_k} - \frac{M_k C^T}{r_k} L_k^T + L_k L_k^T\right)$$
$$= M_k + r_k \left(\left(L_k - \frac{M_k C^T}{r_k}\right)\left(L_k - \frac{M_k C^T}{r_k}\right)^T - \frac{M_k C^T}{r_k}\frac{C M_k}{r_k}\right)$$

The value of $L_k$ which minimises $P_{k+1}$ is therefore given by:

$$L_k^{opt} = \frac{M_k C^T}{r_k} = \frac{M_k C^T}{C M_k C^T + p_{std}^2} \tag{8}$$

The optimal estimation error is then:

$$P_{k+1}^{opt} = M_k - \frac{M_k C^T C M_k}{r_k} = M_k - \frac{M_k C^T C M_k}{C M_k C^T + p_{std}^2} \tag{9}$$

The optimal prediction error is obtained by injecting equation 9 into equation 6:

$$M_{k+1}^{opt} = A P_{k+1}^{opt} A^T + a_{std}^2 B B^T = A \left( M_k - \frac{M_k C^T C M_k}{C M_k C^T + p_{std}^2} \right) A^T + a_{std}^2 B B^T \tag{10}$$

### Steady state

We assume that the recursive equations for the optimal gain (equation 8), estimation error (equation 9) and prediction error (equation 10) have converged to their steady-state values. We wish to determine the steady-state optimal gain, which we write:

$$L = \begin{pmatrix} l_1 \\ \frac{l_2}{T} \end{pmatrix} \tag{11}$$

This can be obtained from the steady state prediction error which we write:

$$M = \begin{pmatrix} m_1 & \frac{m_2}{T} \\ \frac{m_2}{T} & \frac{m_3}{T^2} \end{pmatrix}$$

Indeed, using equation 8, we have:

$$l_1 = \frac{m_1}{m_1 + p_{std}^2} \tag{12}$$

$$l_2 = \frac{m_2}{m_1 + p_{std}^2} \tag{13}$$

We start by determining a system of three equations with three unkowns $m_1, m_2, m_3$. Replacing with equations 12 and 13, we then determine a system of two equations with unkowns $l_{1,2}$, which we solve to determine the optimal steady-state gains.

#### Steady state prediction error

The steady state prediction error $M$ is a fixed point of the recursive equation 10. Therefore, it verifies:

$$
\begin{aligned}
M &= A\left(M - \frac{MC^TCM}{CMC^T + p_{std}^2}\right)A^T + a_{std}^2 BB^T \\
0 &= -M + AMA^T - \frac{AMC^TCMA^T}{CMC^T + p_{std}^2} + a_{std}^2 BB^T \qquad (14)
\end{aligned}
$$

We write:

Equation 14 corresponds to a system of three equations with three unkowns $m_1, m_2, m_3$. We start by writing out these three equations explicitly.

We have:

$$
MA^T = \begin{pmatrix} m_1 & \frac{m_2}{T} \\ \frac{m_2}{T} & \frac{m_3}{T^2} \end{pmatrix} \begin{pmatrix} 1 & 0 \\ T & 1 \end{pmatrix} = \begin{pmatrix} m_1 + m_2 & \frac{m_2}{T} \\ \frac{m_2+m_3}{T} & \frac{m_3}{T^2} \end{pmatrix}
$$

Therefore:

$$
AMA^T = \begin{pmatrix} 1 & T \\ 0 & 1 \end{pmatrix} \begin{pmatrix} m_1 + m_2 & \frac{m_2}{T} \\ \frac{m_2+m_3}{T} & \frac{m_3}{T^2} \end{pmatrix} = \begin{pmatrix} m_1 + 2m_2 + m_3 & \frac{m_2+m_3}{T} \\ \frac{m_2+m_3}{T} & \frac{m_3}{T^2} \end{pmatrix}
$$

$$
-M + AMA^T = \begin{pmatrix} 2m_2 + m_3 & \frac{m_3}{T} \\ \frac{m_3}{T} & 0 \end{pmatrix}
$$

Moreover:

$$
CMA^T = \begin{pmatrix} 1 & 0 \end{pmatrix} \begin{pmatrix} m_1 + m_2 & \frac{m_2}{T} \\ \frac{m_2+m_3}{T} & \frac{m_3}{T^2} \end{pmatrix} = \begin{pmatrix} m_1 + m_2 & \frac{m_2}{T} \end{pmatrix}
$$

Therefore:

$$
\begin{aligned}
AMC^TCMA^T = (CMA^T)^T CMA^T &= \begin{pmatrix} m_1 + m_2 \\ \frac{m_2}{T} \end{pmatrix} \begin{pmatrix} m_1 + m_2 & \frac{m_2}{T} \end{pmatrix} \\
&= \begin{pmatrix} (m_1 + m_2)^2 & (m_1 + m_2)\frac{m_2}{T} \\ (m_1 + m_2)\frac{m_2}{T} & \frac{m_2^2}{T^2} \end{pmatrix}
\end{aligned}
$$

Finally,

$$
BB^T = T^4 \begin{pmatrix} \frac{1}{2} \\ \frac{1}{T} \end{pmatrix} \begin{pmatrix} \frac{1}{2} & \frac{1}{T} \end{pmatrix} = T^4 \begin{pmatrix} \frac{1}{4} & \frac{1}{2T} \\ \frac{1}{2T} & \frac{1}{T^2} \end{pmatrix}
$$

Replacing all the above into equation 14 and using $CMC^T = m_1$, we obtain:

$$
0 = \begin{pmatrix} 2m_2 + m_3 & \frac{m_3}{T} \\ \frac{m_3}{T} & 0 \end{pmatrix} - \frac{1}{m_1 + p_{std}^2} \begin{pmatrix} (m_1 + m_2)^2 & (m_1 + m_2)\frac{m_2}{T} \\ (m_1 + m_2)\frac{m_2}{T} & \frac{m_2^2}{T^2} \end{pmatrix} + (a_{std}T^2)^2 \begin{pmatrix} \frac{1}{4} & \frac{1}{2T} \\ \frac{1}{2T} & \frac{1}{T^2} \end{pmatrix}
$$

This gives the following system of three equations with three unkowns:

$$0 = 2m_2 + m_3 - \frac{(m_1+m_2)^2}{m_1+p_{std}^2} + \frac{(a_{std}T^2)^2}{4} \quad (15)$$

$$0 = m_3 - \frac{(m_1+m_2)m_2}{m_1+p_{std}^2} + \frac{(a_{std}T^2)^2}{2} \quad (16)$$

$$0 = 0 - \frac{m_2^2}{m_1+p_{std}^2} + (a_{std}T^2)^2 \Rightarrow (a_{std}T^2)^2 = \frac{m_2^2}{m_1+p_{std}^2} \quad (17)$$

#### Steady state gain

To obtain a system of two equations with unkowns $l_1, l_2$, we start by removing $m_3$ from the above equations by subtracting equation 16 from equation 15:

$$\begin{aligned} 0 &= 2m_2 + \frac{(m_1+m_2)}{m_1+p_{std}^2}(m_2 - m_1 - m_2) - \frac{(a_{std}T^2)^2}{4} \\ &= 2m_2 - m_1\frac{(m_1+m_2)}{m_1+p_{std}^2} - \frac{(a_{std}T^2)^2}{4} \end{aligned} \quad (18)$$

Replacing equation 17 into equation 18:

$$2m_2 - m_1\frac{(m_1+m_2)}{m_1+p_{std}^2} - \frac{1}{4}\frac{m_2^2}{m_1+p_{std}^2} \quad (19)$$

Dividing by $m_1 + p_{std}^2$ we can replace $m_1$ and $m_2$ with the steady state gains $l_1$ and $l_2$ using equations 12 and 13:

$$0 = 2l_2 - l_1(l_1 + l_2) - \frac{1}{4}l_2^2 \quad (20)$$

We now need to express $l_1$ as a function of $l_2$. Dividing equation 17 by $m_1 + p_{std}^2$ and replacing with the equation for $l_2$ (equation 13):

$$l_2^2 = \frac{(a_{std}T^2)^2}{m_1+p_{std}^2} \quad (21)$$

Using equation 12, we have:

$$\begin{aligned} m_1 = l_1(m_1+p_{std}^2) \Rightarrow m_1(1-l_1) = l_1p_{std}^2 \Rightarrow m_1 = p_{std}^2\frac{l_1}{1-l_1} \\ \Rightarrow m_1 + p_{std}^2 = p_{std}^2(1 + \frac{l_1}{1-l_1}) = p_{std}^2\frac{1-l_1+l_1}{1-l_1} = \frac{p_{std}^2}{1-l_1} \end{aligned} \quad (22)$$

Using equation 18, we can remove $m_1$ from equation 21:

$$l_2^2 = \frac{(a_{std}T^2)^2}{p_{std}^2}(1-l_1) \quad (23)$$

Introducing the dimensionless ratio $r = \frac{p_{std}}{a_{std}T^2}$, equation 23 can be rewritten as:

$$r^2 l_2^2 = 1 - l_1$$
$$l_1 = 1 - r^2 l_2^2 \quad (24)$$

Using equation 24, we can remove $l_1$ from equation 20:

$$\begin{aligned} 0 &= 2l_2 - (1 - r^2l_2^2)(1 - r^2l_2^2 + l_2) - \frac{1}{4}l_2^2 \\ &= 2l_2 - 1 + r^2l_2^2 - l_2 + r^2l_2^2 - (r^2l_2^2)^2 + r^2l_2^3 - \frac{1}{4}l_2^2 \\ 0 &= r^4l_2^4 - r^2l_2^3 + l_2^2(\frac{1}{4} - 2r^2) - l_2 + 1 \end{aligned} \quad (25)$$

Multiplying by $r^4$, we find that $x = r^2 l_2$ verifies:

$$0 = x^4 - x^3 + x^2\left(\frac{1}{4} - 2r^2\right) - r^2x + r^4 \quad (26)$$

This polynomial has four roots. To determine the steady-state gain, we must determine the root $x^*$ for which both $l_2 = x^*/r^2$ and $l_1 = 1 - r^2l_2^2 = 1 - (x^*)^2/r^2$ are real and positive over the whole range of $r$. We start by decomposing the fourth order polynomial in equation 26 into a product of two second order polynomials:

$$x^4 - x^3 + x^2\left(\frac{1}{4} - 2r^2\right) - r^2x + r^4 = \left(x^2 + x\left(-\frac{1}{2} + 2r\right) + r^2\right)\left(x^2 - x\left(\frac{1}{2} + 2r\right) + r^2\right)$$

The first of these second order polynomials has discriminant:

$$\Delta_1 = \left(-\frac{1}{2} + 2r\right)^2 - 4r^2 = \frac{1}{4} - 2r$$

For $r > \frac{1}{8}$, we have $\Delta_1 < 0$ and this first second order polynomial has two complex roots. Therefore, the roots of this polynomial do not correspond to $x^*$.

Next, we determine the roots of the polynomial $x^2 - x\left(\frac{1}{2} + 2r\right) + r^2$. Its discriminant is:

$$\Delta_2 = \left(\frac{1}{2} + 2r\right)^2 - 4r^2 = \frac{1}{4} + 2r > 0$$

Its roots are:

$$x_\pm = \frac{1}{2}\left(\left(\frac{1}{2}+2r\right) \pm \sqrt{\frac{1}{4}+2r}\right) = \frac{1}{4}+r \pm \frac{1}{4}\sqrt{1+8r}$$

To determine which of these roots corresponds to $x^*$ , we calculate $1 - x_\pm^2/r^2$, which must be positive over the whole range of $r$.

$$\begin{aligned}
1-\frac{x_+^2}{r^2} &= 1-\frac{1}{r^2}\left(\frac{1}{16}+r^2+\frac{1+8r}{16}+\frac{r}{2}+\frac{1}{8}\sqrt{1+8r}+r\frac{1}{2}\sqrt{1+8r}\right)\\
&= -\frac{1}{r^2}\left(-r^2+\frac{1}{16}+r^2+\frac{1+8r}{16}+\frac{r}{2}+\frac{1}{8}\sqrt{1+8r}+r\frac{1}{2}\sqrt{1+8r}\right)\\
&= -\frac{1}{r^2}\left(\frac{1}{16}+\frac{1+8r}{16}+\frac{r}{2}+\frac{1}{8}\sqrt{1+8r}+r\frac{1}{2}\sqrt{1+8r}\right) < 0
\end{aligned}$$

Therefore $x_+$ does not correspond to $x^*$.

$$\begin{aligned}
1-\frac{x_-^2}{r^2} &= 1-\frac{1}{r^2}\left(\frac{1}{16}+r^2+\frac{1+8r}{16}+\frac{r}{2}-\frac{1}{8}\sqrt{1+8r}-\frac{r}{2}\sqrt{1+8r}\right)\\
&= -\frac{1}{r^2}\left(\frac{1}{16}+\frac{1+8r}{16}+\frac{r}{2}-\frac{1}{8}\sqrt{1+8r}-\frac{r}{2}\sqrt{1+8r}\right)\\
&= -\frac{1}{r^2}\left(\frac{1}{8}+r-(\frac{1}{8}+\frac{r}{2})\sqrt{1+8r}\right) = -\frac{1}{8r^2}\left(1+8r-(1+4r)\sqrt{1+8r}\right)\\
&= -\frac{1}{r^2}\sqrt{1+8r}\left(\sqrt{1+8r}-1-4r\right)
\end{aligned}$$

$$1-\frac{x_-^2}{r^2} > 0 \Leftrightarrow 1+4r > \sqrt{1+8r} \Leftrightarrow 1+8r+16r^2 > 1+8r \Leftrightarrow r^2 > 0$$

Therefore, $x_-$ corresponds to $x^*$ and the optimal gains are given by (Figure 1):

$$\begin{aligned}
l_2 &= \frac{x_-}{r^2} = \frac{1+4r-\sqrt{1+8r}}{4r^2} &\quad (27)\\
l_1 &= 1-r^2l_2^2 = \frac{\left(\sqrt{1+8r}\right)(1+4r)-1-8r}{8r^2} &\quad (28)
\end{aligned}$$

##### Small $p_{std}$ and small $a_{std}$ limits

In the limit where $p_{std} \to 0$, we have $r \to 0$. We do a Taylor expansion:

$$\begin{aligned}
l_2 &= \frac{1+4r-\sqrt{1+8r}}{4r^2} \approx \frac{1+4r-(1+4r-(8r)^2/8)}{4r^2} = 2\\
l_1 &= 1-r^2l_2^2 \approx 1
\end{aligned}$$

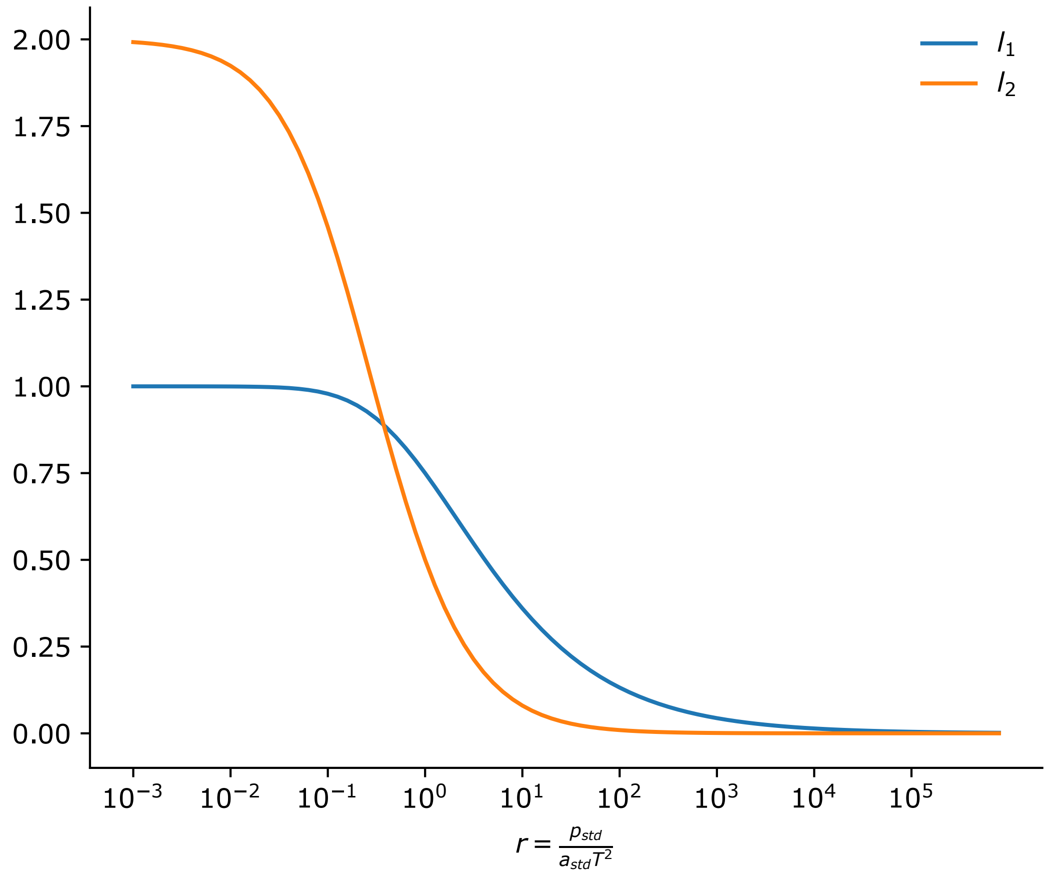

Figure 1: Estimator gains as a function of the ratio of position to acceleration noise

Note that, in this limit, $l_1 = 1$, therefore the position estimate exactly follows the position measurement.

In the limit where $a_{std} \to 0$, we have $r \to +\inf$.

$$\begin{aligned} l_2 &= \frac{1 + 4r - \sqrt{1+8r}}{4r^2} \approx \frac{4r}{4r^2} = \frac{1}{r} \to 0 \\ l_1 &= 1 - r^2 l_2^2 \to 0 \end{aligned}$$

Note that, in this limit, $l_1 = l_2 = 0$, therefore the position measurement is ignored and the position estimate is obtained by double integration of the acceleration measurement.

### Estimator transfer function

The estimate is a linear combination of the position and acceleration measurements. To gain insight into how these measurements are combined, we calculate the steady-state transfer function from from position and acceleration measurements to the estimate.

#### Discrete Fourier transform

We introduce the discrete Fourier transform: the Fourier coefficients (which we denote by a tilde) of a finite sequence $\{s_{n=0,...,N-1}\}$ of length $N$ are given by:

$$\forall j \in [0, ..., N-1], \tilde{s}_j = \sum_{k=0}^{N-1} s_k \exp\left(-i\frac{2\pi jk}{N}\right)$$

The original sequence $\{s_{n=0,...,N-1}\}$ can then be obtained from the Fourier coefficients using the inverse discrete Fourier transform:

$$\begin{aligned} \forall k \in [0, ..., N-1], s_k &= \frac{1}{N}\sum_{j=0}^{N-1} \tilde{s}_j \exp\left(i\frac{2\pi jk}{N}\right) = \frac{1}{N}\sum_{j=0}^{N-1} \tilde{s}_j \left(\exp\left(i\frac{2\pi j}{N}\right)\right)^k \\ &= \frac{1}{N}\sum_{j=0}^{N-1} \tilde{s}_j (z_j)^k \end{aligned}$$

where $z_j = \exp\left(-i\frac{2\pi j}{N}\right)$.

We introduce the discrete Fourier coefficients $\tilde{x}_j^{meas}, \tilde{a}_j^{meas}, \tilde{X}_j^{est}$ of the position measurement, acceleration measurement and estimate. The original signals can then be obtained using the inverse discrete Fourier transform:

$$\forall k \in [0, ..., N-1], a_k^{meas} = \frac{1}{N} \sum_{j=0}^{N-1} \tilde{a}_j^{meas} (z_j)^k \tag{29}$$

$$x_k^{meas} = \frac{1}{N} \sum_{j=0}^{N-1} \tilde{x}_j^{meas} (z_j)^k \tag{30}$$

$$X_k^{est} = \frac{1}{N} \sum_{j=0}^{N-1} \tilde{X}_j^{est} (z_j)^k \tag{31}$$

#### Discrete Fourier transform of the estimator dynamics

To obtain the transfer functions from position and acceleration measurements to the estimate, we need to express $\tilde{X}_j^{est}$ as a linear combination of $\tilde{x}_j^{meas}$ and $\tilde{a}_j^{meas}$. For this, we first express the steady-state estimator dynamics in terms of the position and acceleration measurements, by replacing equation 2 into equation 4:

$$X_{k+1}^{est} = (1 - LC)(AX_k^{est} + Ba_k^{meas}) + Lx_{k+1}^{meas} \tag{32}$$

To simplify notations, we introduce:

$$D = (1 - LC)A$$
$$U = (1 - LC)B$$

Replacing in equation 32, we obtain:

$$X_{k+1}^{est} = DX_k^{est} + Ua_k^{meas} + Lx_{k+1}^{meas}$$
$$\Rightarrow X_{k+1}^{est} - DX_k^{est} = Ua_k^{meas} + Lx_{k+1}^{meas} \tag{33}$$

We now replace the expressions in equations 29 to 31 into equation 33, using:

$$x_{k+1}^{meas} = \frac{1}{N} \sum_{j=0}^{N-1} \tilde{x}_j^{meas} (z_j)^{k+1} = \frac{1}{N} \sum_{j=0}^{N-1} \tilde{x}_j^{meas} (z_j)^k z_j$$

$$X_{k+1}^{est} = \frac{1}{N} \sum_{j=0}^{N-1} \tilde{X}_j^{est} (z_j)^k z_j$$

$$X_{k+1}^{est} - DX_k^{est} = Ua_k^{meas} + Lx_{k+1}^{meas}$$

$$= \frac{1}{N} \sum_{j=0}^{N-1} (z_j)^k (z_j - D) \tilde{X}_j^{est} = \frac{1}{N} \sum_{j=0}^{N-1} (z_j)^k \left( U\tilde{a}_j^{meas} + z_j L \tilde{x}_j^{meas} \right) \tag{34}$$

For equation 34 to be valid for all $k$ in $[0, ..., N-1]$, we must have:

$$\forall j \in [0, ..., N-1], (z_j - D)\tilde{X}_j^{est} = \left( U\tilde{a}_j^{meas} + z_j L \tilde{x}_j^{meas} \right)$$

We can thus express $\tilde{X}_j^{est}$ as a function of $\tilde{a}_j^{meas}$ and $\tilde{x}_j^{meas}$:

$$\begin{aligned}\forall j \in [0,...,N-1], \tilde{X}_j^{est} &= (z_j - D)^{-1} U \tilde{a}_j^{meas} + z_j (z_j - D)^{-1} L \tilde{x}_j^{meas} \\ &= H_{a\to X}(z_j)\tilde{a}_j^{meas} + H_{x\to X}(z_j)\tilde{x}_j^{meas}\end{aligned}$$

where $H_{a\to X} : z \to (z-D)^{-1} U$ is the transfer function from acceleration measurement to estimate, and $H_{x\to X} : z \to z\,(z-D)^{-1} L$ is the transfer function from position measurement to estimate.

#### Calculation of the estimator transfer functions

To obtain explicit expressions for $H_{a\to X}(z)$ and $H_{x\to X}(z)$, we start by showing that:

$$(z_j - D)^{-1} = \frac{1}{\det(z_j - D)}(z_j - \det(D)D^{-1}) \tag{35}$$

For this, we write:

$$D = \begin{pmatrix} d_{11} & d_{12} \\ d_{21} & d_{22} \end{pmatrix}$$

We use the fact that the inverse of any such 2 by 2 matrix $D$ is given by:

$$D^{-1} = \frac{1}{\det(D)} \begin{pmatrix} d_{22} & -d_{12} \\ -d_{21} & d_{11} \end{pmatrix}$$

Indeed:

$$\begin{pmatrix} d_{22} & -d_{12} \\ -d_{21} & d_{11} \end{pmatrix} \begin{pmatrix} d_{11} & d_{12} \\ d_{21} & d_{22} \end{pmatrix} = \begin{pmatrix} d_{22}d_{11} - d_{12}d_{21} & 0 \\ 0 & d_{12}d_{21} + d_{22}d_{11} \end{pmatrix} = \det(D) \begin{pmatrix} 1 & 0 \\ 0 & 1 \end{pmatrix}$$

We likewise have:

$$\begin{aligned}(z_j - D)^{-1} &= \frac{1}{\det(z_j - D)} \begin{pmatrix} z_j - d_{22} & d_{12} \\ d_{21} & z_j - d_{11} \end{pmatrix} \\ &= \frac{1}{\det(z_j - D)}(z_j - \det(D)D^{-1})\end{aligned}$$

To obtain an expression for $\det(D)$, we write:

$$\det(D) = \det\left((1 - LC)A\right) = \det(1 - LC)\det(A) = \det(1 - LC)$$

$$1 - LC = 1 - \begin{pmatrix} l_1 \\ \frac{l_2}{T} \end{pmatrix} \begin{pmatrix} 1 & 0 \end{pmatrix} = \begin{pmatrix} 1 - l_1 & 0 \\ -\frac{l_2}{T} & 1 \end{pmatrix} \tag{36}$$

$$\Rightarrow \det(D) = \det(1 - LC) = 1 - l_1 \tag{37}$$

To obtain an expression for $\det(z_j - D)$, we write:

$$\det(z_j - D) = (z_j - d_{11})(z_j - d_{22}) - d_{21}d_{12} = z_j^2 - z_j(d_{11} + d_{22}) + \det(D)$$

To obtain an expression for $d_{11} + d_{22}$, we explicitly calculate $D$ (using equation 36):

$$D = (1 - LC)A = \begin{pmatrix} 1 - l_1 & 0 \\ -\frac{l_2}{T} & 1 \end{pmatrix} \begin{pmatrix} 1 & T \\ 0 & 1 \end{pmatrix} = \begin{pmatrix} 1 - l_1 & T(1 - l_1) \\ -\frac{l_2}{T} & 1 - l_2 \end{pmatrix}$$

$$\Rightarrow d_{11} + d_{22} = 2 - (l_1 + l_2)$$

$$\Rightarrow \det(z_j - D) = z_j^2 - z_j \left(2 - (l_1 + l_2)\right) + 1 - l_1 \qquad (38)$$

**Transfer function from the position measurement**

The transfer function from the position measurement is given by:

$$\begin{aligned} H_{x \to X}(z_j) &= (z_j - D)^{-1} L = \frac{1}{\det(z_j - D)} (z_j - \det(D) D^{-1}) L \\ &= \frac{1}{\det(z_j - D)} (z_j - \det(D) A^{-1} (1 - LC)^{-1}) L \end{aligned} \qquad (39)$$

We have:

$$(1-LC)^{-1} L = \frac{1}{\det(1 - LC)} \begin{pmatrix} 1 & 0 \\ \frac{l_2}{T} & 1 - l_1 \end{pmatrix} \begin{pmatrix} l_1 \\ \frac{l_2}{T} \end{pmatrix} = \frac{1}{\det(D)} \begin{pmatrix} l_1 \\ \frac{l_2}{T}(l_1 + 1 - l_1) \end{pmatrix} = \frac{1}{\det(D)} L$$

Reinjecting this into equation 39, we have:

$$\begin{aligned} H_{x \to X}(z_j) &= \frac{1}{\det(z_j - D)} (z_j - A^{-1}) L = \frac{1}{\det(z_j - D)} \begin{pmatrix} z_j - 1 & T \\ 0 & z_j - 1 \end{pmatrix} \begin{pmatrix} l_1 \\ \frac{l_2}{T} \end{pmatrix} \\ &= \frac{1}{\det(z_j - D)} \begin{pmatrix} l_1(z_j - 1) + l_2 \\ \frac{l_2}{T}(z_j - 1) \end{pmatrix} \end{aligned} \qquad (40)$$

Injecting equation 38 into equation 40, we obtain the transfer function from the position measurement to the position estimate $x^{est}$ (Figure 2.A):

$$H_{x \to x^{est}}(z_j) = \frac{l_1(z_j - 1) + l_2}{z_j^2 - z_j \left(2 - (l_1 + l_2)\right) + 1 - l_1} \qquad (41)$$

Likewise, the transfer function from the position measurement to the velocity estimate $v^{est}$ is given by:

$$H_{x \to v^{est}}(z_j) = \frac{1}{T} \frac{l_2(z_j - 1)}{z_j^2 - z_j \left(2 - (l_1 + l_2)\right) + 1 - l_1}$$

##### Transfer function from the acceleration measurement

The transfer function from the acceleration measurement is given by:

$$\begin{aligned}
H_{x\to X}(z_j) &= (z_j - D)^{-1}U = \frac{1}{\det(z_j - D)}(z_j - \det(D)D^{-1})(1 - LC)B \\
&= \frac{1}{\det(z_j - D)}\left(z_j(1 - LC) - \det(D)A^{-1}(1 - LC)^{-1}(1 - LC)\right)B \\
&= \frac{1}{\det(z_j - D)}\left(z_j\begin{pmatrix}1 - l_1 & 0\\ -\frac{l_2}{T} & 1\end{pmatrix} - (1 - l_1)\begin{pmatrix}1 & -T\\ 0 & 1\end{pmatrix}\right)B \\
&= \frac{1}{\det(z_j - D)}\begin{pmatrix}(z_j - 1)(1 - l_1) & T(1 - l_1)\\ -z_j\frac{l_2}{T} & z_j - (1 - l_1)\end{pmatrix}\begin{pmatrix}\frac{T^2}{2}\\ T\end{pmatrix} \\
&= \frac{1}{\det(z_j - D)}\begin{pmatrix}\frac{T^2}{2}(1 - l_1)(z_j - 1 + 2)\\ T(-\frac{l_2}{2}z_j + z_j - 1 + l_1)\end{pmatrix} \\
&= \frac{1}{\det(z_j - D)}\begin{pmatrix}\frac{T^2}{2}(1 - l_1)(z_j + 1)\\ T(z_j(1 - \frac{l_2}{2}) - 1 + l_1)\end{pmatrix} \qquad (42)
\end{aligned}$$

Injecting equation 38 into equation 42, we obtain the transfer function from the acceleration measurement to the position estimate:

$$H_{a\to x^{est}}(z_j) = \frac{T^2}{2}\frac{(1 - l_1)(z_j + 1)}{z_j^2 - z_j\left(2 - (l_1 + l_2)\right) + 1 - l_1} \qquad (43)$$

Likewise, the transfer function from the acceleration measurement to the velocity estimate is given by:

$$H_{a\to v^{est}}(z_j) = T\frac{z_j(1 - \frac{l_2}{2}) - 1 + l_1}{z_j^2 - z_j\left(2 - (l_1 + l_2)\right) + 1 - l_1}$$

##### Transfer function of double integration

We wish to compare the estimator to the state estimate $X^{int}$ obtained by integration of the acceleration measurement. This state estimate follows the dynamics:

$$X_{k+1}^{int} = AX_k^{int} + Ba_k^{meas}$$

A derivation similar to the one presented above yields the following transfer function from the acceleration measurement to the integrated state:

$$\begin{aligned}
H_{a\to X^{int}}(z_j) &= (z_j - A)^{-1}B = \frac{1}{\det(z_j - A)}(z - \det(A)A^{-1})B \\
&= \frac{1}{\det(z_j - A)}\begin{pmatrix}z_j - 1 & T\\ 0 & z_j - 1\end{pmatrix}\begin{pmatrix}\frac{T^2}{2}\\ T\end{pmatrix} = \frac{1}{\det(z_j - A)}\begin{pmatrix}\frac{T^2}{2}(z_j + 1)\\ T(z_j - 1)\end{pmatrix} \\
\det(z_j - A) &= \det\begin{pmatrix}z_j - 1 & -T\\ 0 & z_j - 1\end{pmatrix} = (z_j - 1)^2
\end{aligned}$$

The transfer function from the acceleration measurement to the position obtained by integration $x^{int}$ is thus:

$$H_{a\rightarrow x^{int}}(z_j) = \frac{T^2}{2}\frac{z_j+1}{(z_j-1)^2}$$

The ratio of the estimator and integrator positions (Figure 2.B) is given by:

$$\begin{aligned}\frac{H_{a\rightarrow x^{est}}(z_j)}{H_{a\rightarrow x^{int}}(z_j)} &= (1-l_1)\frac{(z_j-1)^2}{z_j^2-z_j\,(2-(l_1+l_2))+1-l_1} \\ &= (1-l_1)\frac{(z_j-1)^2}{(z_j-1)^2+z_j(l_1+l_2)-l_1} \qquad (44)\end{aligned}$$

Note that, at low frequencies (i.e. when $z_j \rightarrow 1$), the integrator transfer function $H_{a\rightarrow x^{int}}(z_j)$ diverges whereas the estimator transfer function $H_{a\rightarrow x^{est}}(z_j)$ reaches a finite value $\frac{T^2(1-l_1)^2}{2l_2}$. The ratio of the two thus goes to zero at low frequencies.

The transfer function from the acceleration measurement to the velocity obtained by integration $v^{int}$ is :

$$H_{a\rightarrow v^{int}}(z_j) = \frac{T}{z_j-1}$$

The ratio of the estimator and integrator velocities is given by:

$$\frac{H_{a\rightarrow v^{est}}(z_j)}{H_{a\rightarrow v^{int}}(z_j)} = (z_j-1)\frac{z_j(1-\frac{l_2}{2})-1+l_1}{z_j^2-z_j\,(2-(l_1+l_2))+1-l_1}$$

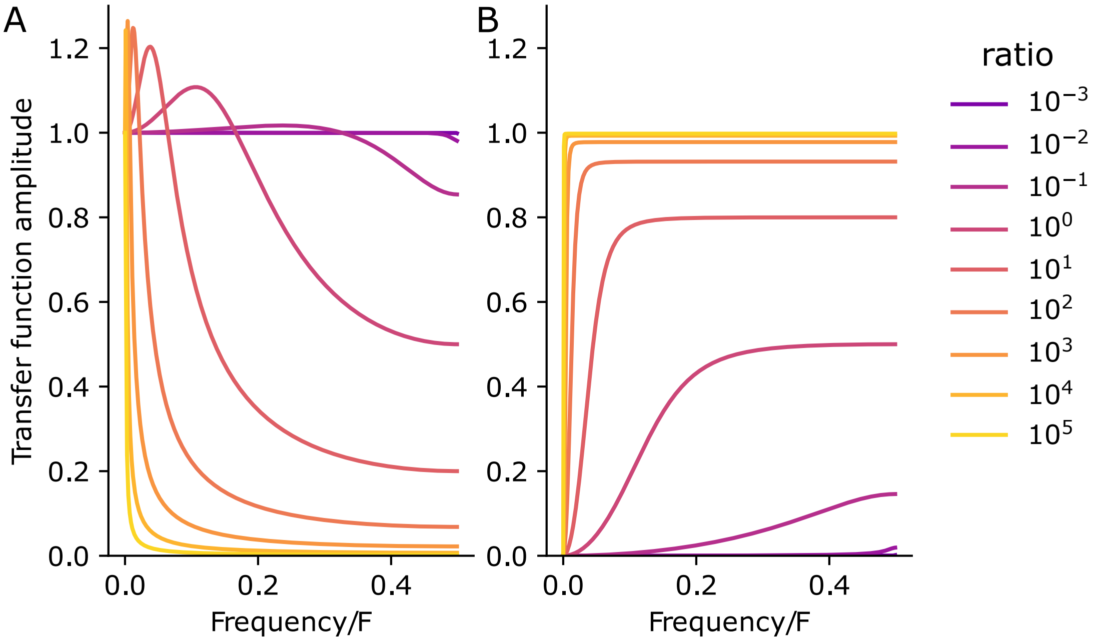

Figure 2: Amplitude of the transfer function (A) from the kinematic to the estimate position, and (B) from the double integral of acceleration to the estimate position, as a function of the frequency divided by the sampling frequency $F = 1/T$, for different values of the noise ratio r
